# Large phospholipid-number asymmetry is not required to reproduce plasma membrane physical properties

**DOI:** 10.64898/2026.09.04.749494

**Authors:** Cristian R. Popov, Marius F.W. Trollmann, Chunzhen Zhang, Rainer A. Böckmann

## Abstract

Plasma membranes are compositionally asymmetric, but whether this lipid-type asymmetry is accompanied by a substantial phospholipid-number imbalance between leaflets remains debated. Here, we use microsecond all-atom molecular dynamics simulations to compare human red blood cell plasma membrane models with either strong phospholipid-number asymmetry and exoplasmic cholesterol enrichment or near-symmetric phospholipid and cholesterol numbers but preserved lipid-type asymmetry. Both asymmetric models reproduce a densely packed, ordered exoplasmic leaflet and a more fluid cytoplasmic leaflet. Strong phospholipid-number asymmetry, however, drives extensive cholesterol enrichment in the exoplasmic leaflet and amplifies membrane asymmetry, leading to large cholesterol-rich clusters, enhanced shallow hydrophobic exposure, reduced exoplasmic lipid mobility, lower ethanol permeability, and increased area compressibility. Comparison with available diffusion and alcohol-permeability measurements indicates that the strongly asymmetric model overestimates the immobilization and barrier properties of the exoplasmic leaflet, whereas the near-symmetric-number model better captures these dynamic observables. Our results suggest that lipid-type asymmetry is sufficient to reproduce many physical hallmarks of plasma membranes without requiring a large phospholipid-number imbalance.

## Introduction

Plasma membranes (PMs) are chemically complex lipid assemblies that define cellular boundaries and regulate membrane protein organization, signaling, trafficking, and membrane remodeling [1–3]. Lipidomic and biochemical studies have shown that PMs exhibit a pronounced transbilayer asymmetry, with distinct lipid compositions in the exoplasmic and cytoplasmic leaflets [4–10]. The exoplasmic leaflet is enriched in saturated phospholipids and sphingomyelin, which promote tight lipid packing and a robust permeability barrier. In contrast, the cytoplasmic leaflet contains larger fractions of unsaturated and anionic phospholipids, providing a more dynamic interface for protein recruitment, membrane remodeling, and signaling [11–14]. This transbilayer asymmetry is essential for membrane mechanics, vesicle trafficking, cytokinesis, apoptosis, membrane fusion, and protein localization. Its regulated disruption, for example through phosphatidylserine exposure, further serves as an important signal in diverse physiological processes [15–26].

In contrast to phospholipid headgroup and lipid tail saturation asymmetry, the transbilayer distribution of cholesterol remains unresolved. Although asymmetric phospholipid organization has been established for decades [4–9], both experimental and computational studies have reached different conclusions as to whether cholesterol is enriched in the exoplasmic leaflet, enriched in the cytoplasmic leaflet, or distributed approximately symmetrically between leaflets [10, 27–32]. The question is further complicated by the possible coupling between cholesterol partitioning and phospholipid number asymmetry. The long-standing assumption that the two leaflets contain comparable numbers of phospholipids [5–7] was recently challenged by Doktorova *et al.* [10], who proposed that the cytoplasmic leaflet of the human red blood cell (huRBC) plasma membrane contains over 50 mol% more phospholipids than the exoplasmic leaflet. Molecular dynamics simulations in the same study, performed for membranes with a 30 mol% phospholipid excess in the cytoplasmic leaflet, showed that phospholipid number asymmetry entails cholesterol redistribution toward the exoplasmic leaflet, resulting in a strongly asymmetric cholesterol distribution. These conclusions have been questioned [33–35], underscoring the need to compare competing huRBC plasma membrane models and to assess their structural and biophysical consequences.

Differences in lipid composition, phospholipid number asymmetry, and cholesterol partitioning are expected to affect membrane physical properties. In particular, cholesterol redistribution and phospholipid asymmetry can modulate lipid packing, bilayer thickness, membrane rigidity, hydrophobic defect formation, permeability, lateral diffusion, curvature stress, and membrane–protein interactions. These properties are central to cellular processes ranging from signaling and transport to membrane remodeling and antimicrobial peptide activity [36–41]. Establishing how alternative PM models differ in these observables is therefore essential for evaluating their physical plausibility and physiological relevance.

Here, we compare atomistic models of the huRBC plasma membrane that differ in phospholipid number asymmetry and cholesterol partitioning (Fig. 1). The first model follows the experimentally derived lipid composition of Lorent *et al.* [9] and contains approximately equal phospholipid numbers and cholesterol contents in the two leaflets. We refer to this model as the symmetric cholesterol distribution (SCD) model. The second model adopts the asymmetric phospholipid distribution proposed by Doktorova *et al.* [10], which is associated with preferential cholesterol accumulation in the exoplasmic leaflet. We refer to this model as the asymmetric cholesterol distribution (ACD) model. As an additional reference, we include a scrambled membrane derived from the SCD model, in which lipid identities are randomized between leaflets while the overall lipid composition is preserved (SCR). Using all-atom molecular dynamics (MD) simulations, we characterize the structural, dynamical, and mechanical characteristics of these membrane models as well as the their permeation properties to determine how alternative assumptions about transbilayer lipid organization shape the physical behavior of the huRBC plasma membrane.

**Fig. 1.**
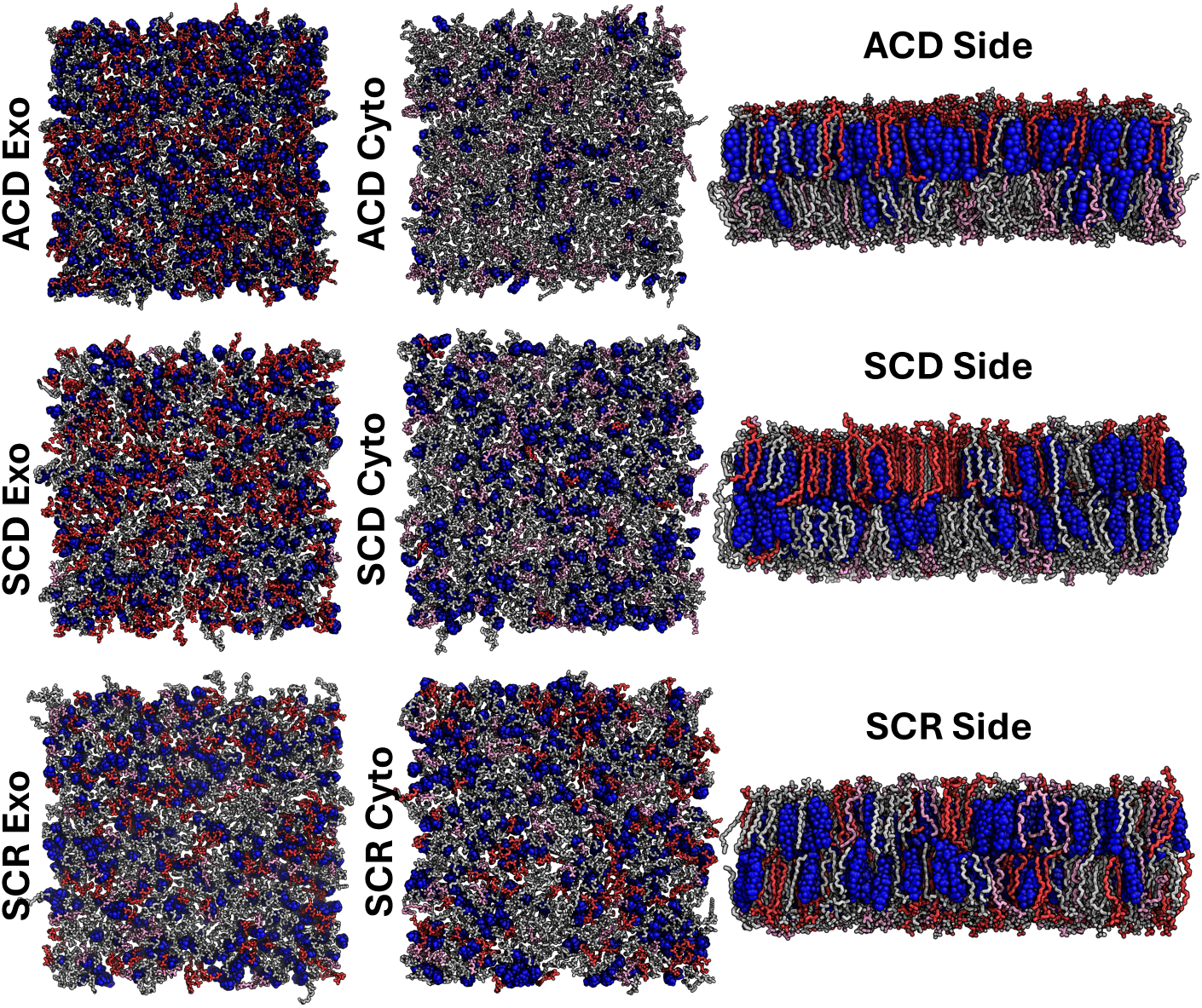
Final-frame snapshots of the ACD, SCD, and SCR membrane models. Exo and Cyto denote the exoplasmic and cytoplasmic leaflets, respectively; Side denotes the membrane side view. Phospholipids are shown as sticks, with sphingomyelins in red, charged lipids such as DOPS and inositides in pink, and all remaining phospholipids in gray. Cholesterol is shown as blue spheres. In the side views, the exoplasmic leaflet is shown on top and the cytoplasmic leaflet on bottom. Images were prepared using PyMOL [45].

## Results

To assess how lipid-type asymmetry and lipid-number asymmetry affect plasma membrane properties, we constructed three all-atom models of the human red blood cell (huRBC) plasma membrane (Fig. 1). The first model contained a 30% excess of phospholipids in the cytoplasmic leaflet and was based on the lipidomics data and simulation-derived leaflet compositions reported by Doktorova *et al.* [10] (ACD model). The second model was based on the lipidomics analysis of Lorent *et al.* [9]. In this model, lipid types were distributed asymmetrically between leaflets, whereas the number of lipids in each leaflet was adjusted according to the lipid areas obtained from simulations of membranes containing either the cytoplasmic or exoplasmic leaflet composition (SCD model). As a reference, we generated a scrambled membrane (SCR), resulting in symmetric lipid-type and lipid-number distributions while preserving the total membrane composition.

All systems contained 40 mol% cholesterol [9, 10]. For the ACD model, the initial asymmetric cholesterol distribution was adopted from Doktorova *et al.* [10]. The membranes were simulated at 310 K and 150 mM NaCl between 13 *µ*s (SCD and SCR) and 20 *µ*s (ACD). The initially (approximately) symmetric cholesterol distributions of the SCD and SCR models, as well as the asymmetric distribution within the ACD model was retained during the course of the simulations.

Cholesterol translocation events between leaflets, hereafter referred to as flip-flops (FFs), were identified and quantified (Fig. S1, Table S1). Cholesterol redistribution was limited in all systems. Net changes amounted to only approximately 1–2% of the total cholesterol population, indicating that the initial cholesterol distributions were largely stable on the simulated timescales. Across the full trajectories, seven successful FF events were observed in both SCD and ACD, whereas five successful events were observed in SCR.

The total number of FF attempts was comparable among the three membrane models. However, the ACD model showed the lowest FF success rate, particularly relative to SCD. This reduced success rate is consistent with the denser packing of the exoplasmic leaflet in ACD, which may hinder cholesterol accommodation after translocation. In line with this interpretation, successful FF events in ACD occurred preferentially from the exoplasmic to the cytoplasmic leaflet. Because only a small number of successful FF events was observed, this directional preference should be interpreted as a qualitative trend rather than a statistically robust difference.

In SCD, cholesterol showed a weak tendency to migrate toward the exoplasmic leaflet. A similar trend was also observed in the symmetric SCR control membrane. Thus, within the present sampling, the net cholesterol flux in SCD cannot be attributed unambiguously to preferential cholesterol interactions with one leaflet composition. Although the number of successful FF events was limited, analysis of the local lipid environment around flip-flopping cholesterol molecules revealed consistent compositional trends (Table S1). Polyunsaturated lipids were enriched around cholesterol molecules undergoing FF, with average enrichment factors of 1.4 ± 0.2 in the initial leaflet and 1.5 ± 0.2 in the final leaflet. In contrast, fully saturated lipids were depleted around cholesterol molecules in FF events, with enrichment factors of 0.6 ± 0.1 and 0.7 ± 0.1 in the initial and final leaflets, respectively. Sterols were also depleted, with corresponding enrichment factors of 0.8 ± 0.1 and 0.9 ± 0.1. Mono- and di-unsaturated lipids showed no strong systematic enrichment or depletion, with enrichment factors of 1.2 ±0.1 and 0.9 ±0.1. These trends suggest that cholesterol translocation is favored in locally more disordered lipid environments and disfavored in tightly packed, saturated, or sterol-rich regions. This interpretation is consistent with previous simulation studies showing facilitated cholesterol flip-flop in more unsaturated membranes and increased flip-flop barriers in cholesterol-rich or raft-like environments [41–44]. Nevertheless, the limited number of observed FF events and the pooling of events across membrane models require cautious interpretation.

### Area per lipid

We first quantified lipid packing by calculating the area per lipid (APL) for each leaflet and membrane model (Fig. 2). In both asymmetric models, ACD and SCD, the exoplasmic leaflet was more densely packed than the cytoplasmic leaflet. This leaflet imbalance was most pronounced in ACD, where the cytoplasmic leaflet APL was 35% larger than the exoplasmic leaflet APL. In SCD, the corresponding difference was only 7% (Fig. 2a). Thus, the phospholipid-number asymmetry in ACD produces a substantially larger packing imbalance between leaflets than the lipid-type asymmetry present in SCD.

**Fig. 2.**
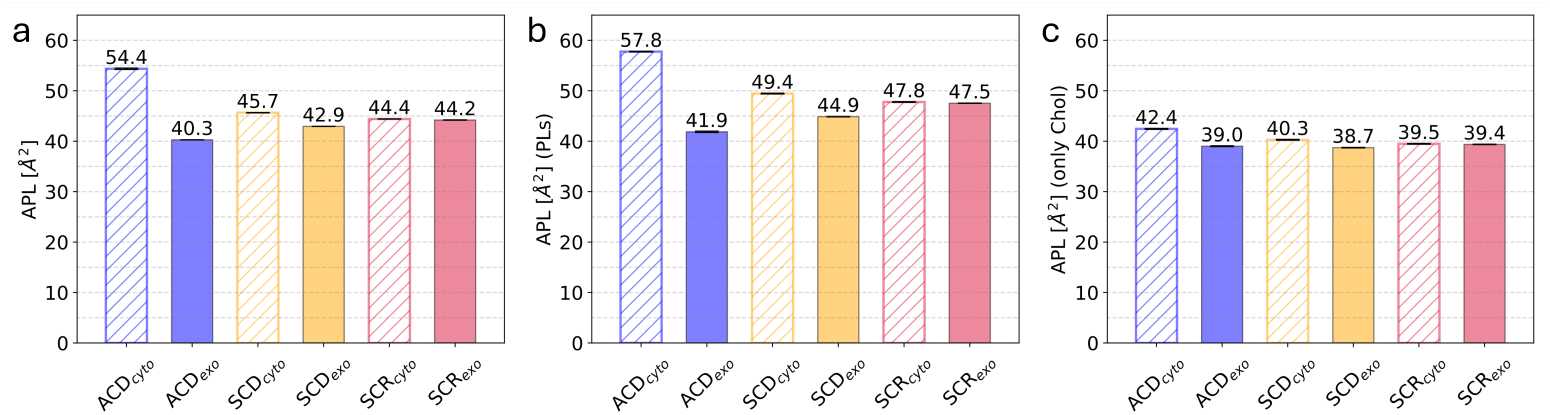
Area per lipid (APL) in the ACD, SCD, and SCR membrane models. (a) Leaflet-averaged APL calculated as the simulation box area divided by the number of lipids, including cholesterol, in the corresponding leaflet. (b,c) Voronoi-based APL calculated from a two-dimensional tessellation of lipid reference atoms for phospholipids and cholesterol. Panel (b) shows phospholipid APL, and panel (c) shows cholesterol APL. Methodological details are provided in Materials and Methods, *Area per Lipid*.

Cholesterol showed only minor differences in Voronoi-based APL among systems and leaflets (Fig. 2c), consistent with its compact and rigid sterol structure. In contrast, phospholipid APL values depended strongly on lipid class and leaflet composition. Sphingomyelins (SMs) occupied smaller areas than phosphatidylcholines (PCs) within the same leaflet (Fig. 3), indicating tighter local packing around SM molecules. This behavior is consistent with favorable cholesterol–SM interactions, which can involve hydrogen bonding between the cholesterol hydroxyl group and the SM sphingosine backbone [46].

**Fig. 3.**
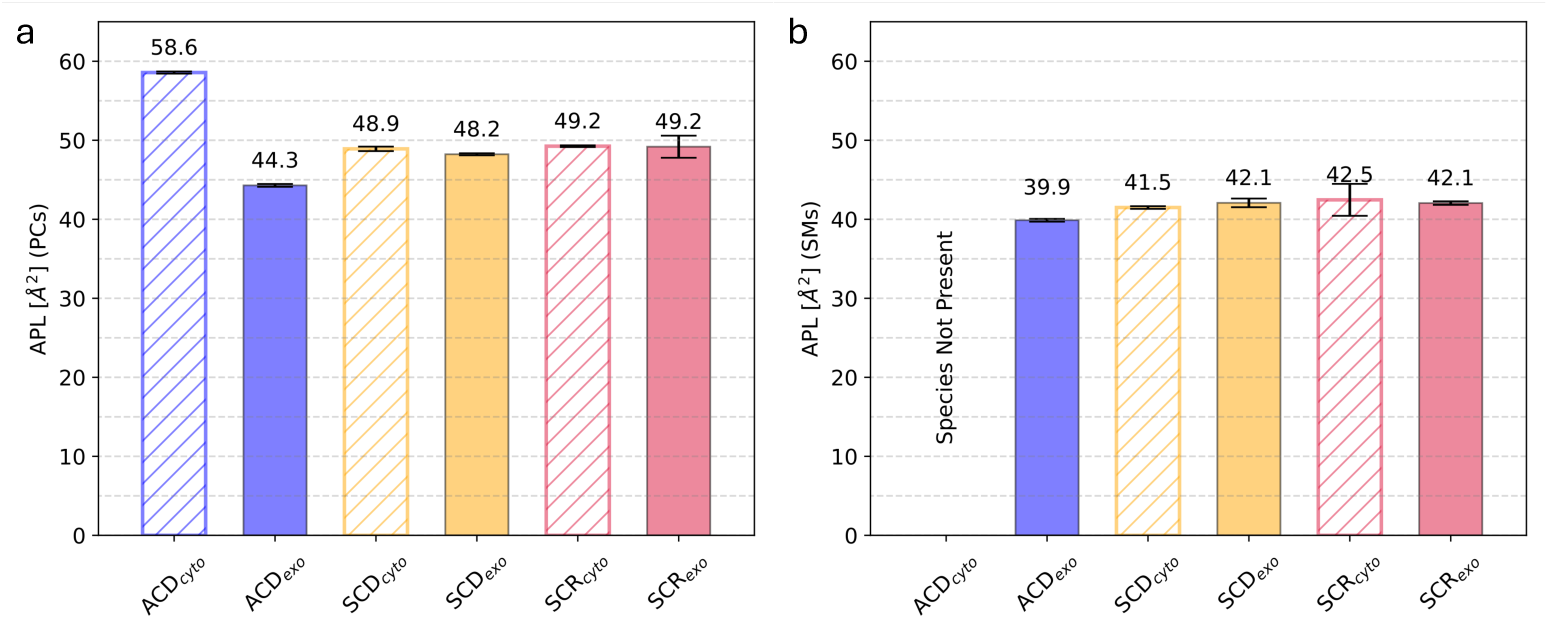
Area per lipid for phosphatidylcholine and sphingomyelin lipids. (a) Average APL of phosphatidylcholine (PC) lipids in their respective leaflets. (b) Average APL of sphingomyelin (SM) lipids in their respective leaflets. APL values were calculated using the same Voronoi-based method as in Fig. 2.

Consistent with tighter cholesterol–SM packing, the cholesterol–PSM radial distribution function showed a first solvation shell at shorter distance than the cholesterol– PLPC radial distribution function (Fig. S4). This geometric difference does not necessarily imply preferential cholesterol–SM association. Indeed, the nearest-neighbor enrichment analysis showed no clear phospholipid preference around cholesterol in the exoplasmic leaflet of ACD (see below). Rather, the radial distribution functions indicate that, when cholesterol contacts SM, these contacts are more compact than corresponding cholesterol–PC contacts.

### Order parameter

The side views in Fig. 1 indicate that the exoplasmic leaflets of the ACD and SCD models are more ordered than their corresponding cytoplasmic leaflets. We quantified this difference using the deuterium order parameter, *S*_CD_, which reports on acyl-chain alignment relative to the membrane normal (see Materials and Methods, *S*_CD_ *Order Parameter*; Fig. 4).

**Fig. 4.**
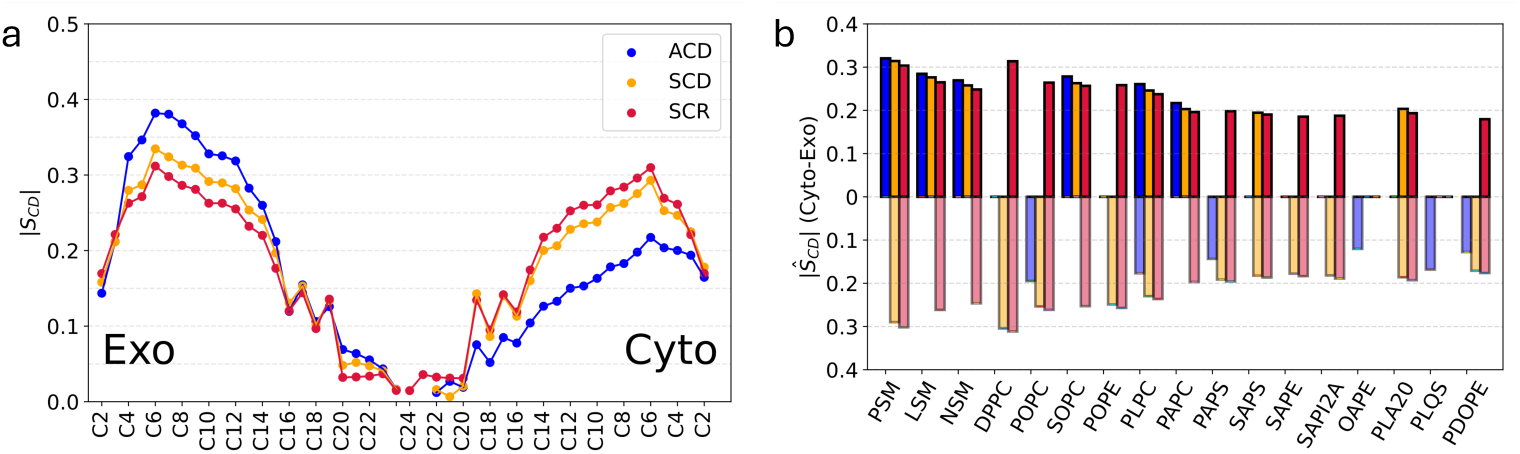
Phospholipid deuterium order parameter. *|S*_CD_*|*. (a) *|S*_CD_*|* profiles resolved by acylchain carbon position relative to the carbonyl region. The center of the plot corresponds to the bilayer midplane, where the two leaflets meet. (b) Lipid-type-averaged order parameters. Semi-transparent bars below the axis correspond to the cytoplasmic leaflet, whereas bars above the axis correspond to the exoplasmic leaflet. Blue, orange, and crimson indicate the ACD, SCD, and SCR plasma membrane models, respectively.

Consistent with the APL analysis, the ACD model showed the strongest leaflet imbalance in lipid packing and order. Its densely packed exoplasmic leaflet was the most ordered leaflet among all models, whereas its cytoplasmic leaflet was the least ordered (Fig. 4). Thus, the phospholipid-number asymmetry in ACD leads to strongly divergent physical properties in the two leaflets.

Across all membrane models, saturated lipids and sphingomyelins exhibited higher acyl-chain order than unsaturated lipid species. Comparison of SCD and SCR further shows that cholesterol density alone does not determine lipid order. These two systems contain the same total cholesterol concentration and retain nearly symmetric cholesterol distributions between leaflets (Table 3). Nevertheless, for the same lipid classes, the exoplasmic leaflet of SCD was more ordered than SCR, whereas the cytoplasmic leaflet of SCD was less ordered than SCR 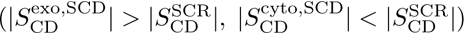. This difference indicates that lipid order is governed not only by cholesterol content but also by leaflet lipid composition.

The most direct compositional determinant is acyl-chain unsaturation. The exoplasmic leaflet of SCD contains approximately 1.2 double bonds per phospholipid, whereas the cytoplasmic leaflet contains approximately 3.4 double bonds per phospholipid. In SCR, the two nearly symmetric leaflets contain approximately 2.3 double bonds per phospholipid (Table 3). The higher order of the SCD exoplasmic leaflet and lower order of the SCD cytoplasmic leaflet therefore follow the expected dependence of acyl-chain order on unsaturation. Together, these data show that leaflet-specific lipid composition, rather than cholesterol density alone, controls the ordering asymmetry of the huRBC plasma membrane models.

### Diffusion

We next analyzed lateral lipid diffusion as a dynamic measure of membrane fluidity, complementary to static descriptors such as the APL (see Materials and Methods, *Lipid Diffusion Coefficients*; Fig. 5). Diffusion coefficients were calculated from the in-plane motion of lipid reference atoms, using phospholipid phosphate atoms and the cholesterol hydroxyl oxygen.

**Fig. 5.**
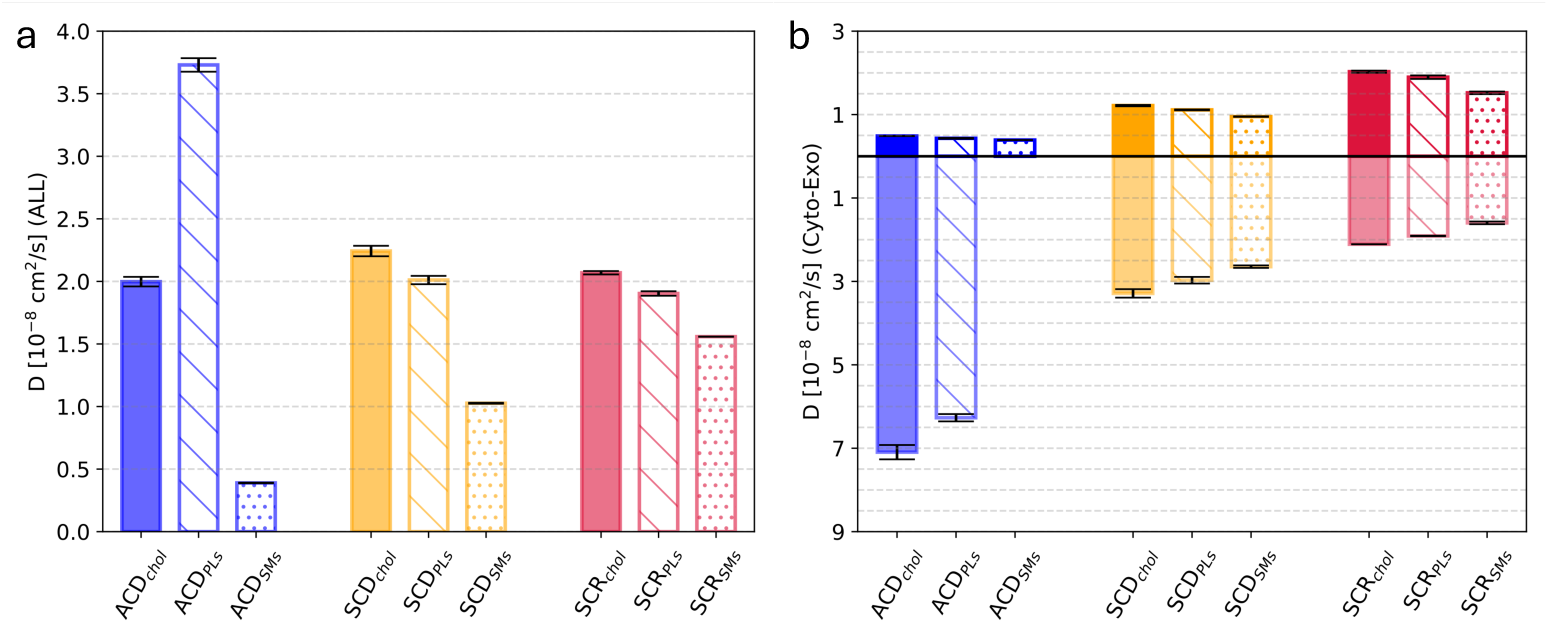
Lipid diffusion coefficients, *D*. (a) Leaflet-averaged diffusion coefficients, weighted by lipid abundance, for cholesterol (chol), phospholipids (PLs), and sphingomyelins (SMs). (b) Corresponding leaflet-resolved diffusion coefficients. Bars below and above the axis correspond to the cytoplasmic and exoplasmic leaflets, respectively. Diffusion was calculated from the in-plane displacement of lipid reference atoms, using phospholipid phosphate atoms and the cholesterol hydroxyl oxygen.

The ACD model showed the strongest leaflet-dependent difference in lateral mobility (Fig. 5b): Lipids in the densely packed exoplasmic leaflet diffused slowly, whereas lipids in the cytoplasmic leaflet were substantially more mobile. This behavior is consistent with the lower APL and higher acyl-chain order of the ACD exoplasmic leaflet, and with the larger APL and lower order of the ACD cytoplasmic leaflet.

Within each leaflet, cholesterol diffused faster than phospholipids in all membrane models (Fig. 5b). The leaflet-averaged ACD data appear to show faster phospholipid diffusion than cholesterol diffusion (Fig. 5a), but this reflects the strongly asymmetric lipid and cholesterol distributions between leaflets. Phospholipids are enriched in the cytoplasmic leaflet of ACD, where they diffuse approximately 12-fold faster than in the exoplasmic leaflet. Cholesterol also diffuses much faster in the cytoplasmic leaflet than in the exoplasmic leaflet, by approximately 14-fold. However, most cholesterol molecules reside in the slowly diffusing exoplasmic leaflet, which lowers the total-averaged cholesterol diffusion coefficient.

Thus, the apparent ranking of lipid mobilities depends on whether diffusion is analyzed in a leaflet-resolved or leaflet-averaged manner. The leaflet-resolved data show that cholesterol is consistently the most mobile lipid class locally, whereas the leaflet-averaged values are strongly shaped by the asymmetric distribution of lipid classes between leaflets. Overall, the diffusion analysis reinforces the conclusion that ACD exhibits the largest transbilayer contrast in physical properties, with a rigid, slowly diffusing exoplasmic leaflet and a comparatively fluid cytoplasmic leaflet.

### Membrane permeability

We next evaluated membrane permeability using ethanol as a molecular probe. Permeation rates of larger solutes, including fluorescein diacetate (FDA) used experimentally [10], are difficult to estimate from all-atom simulations due to their size. For ethanol, we used the accelerated weight histogram (AWH) method, as described in Materials and Methods, *Ethanol Simulations*.

Ethanol experienced a less favorable permeation free-energy landscape in ACD than in SCD (Fig. 6a). This difference is consistent with results obtained for water permeability (Fig. S2) and reflects the higher free-energy barrier encountered by both molecules when crossing the densely packed exoplasmic leaflet of ACD. As a result, the ethanol permeability coefficient was substantially lower in ACD, *P*_EtOH_ = (0.23 ± 0.05) × 10*^−^*^3^ cm*/*s, than in SCD, *P*_EtOH_ = (1.70 ± 0.05) × 10*^−^*^3^ cm*/*s, corresponding to an approximately sevenfold reduction in ethanol permeability in the ACD model.

**Fig. 6.**
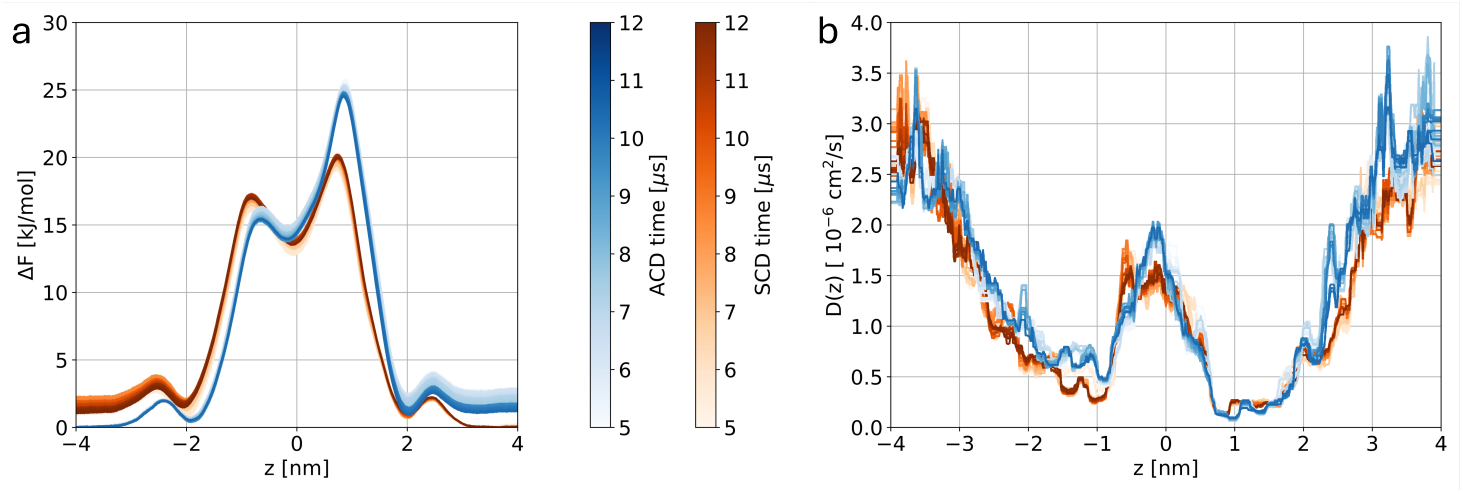
Position-dependent potential of mean force and diffusion coefficient of ethanol. (a) PMF profiles, Δ*F*, for the ACD and SCD plasma membrane models as a function of the distance from the membrane center of mass along the membrane normal, *z*. Negative and positive *z* values correspond to the cytoplasmic and exoplasmic sides, respectively. (b) Position-dependent ethanol diffusion coefficient, *D*(*z*), shown as a 0.2 nm rolling median for the ACD and SCD models. The color gradient indicates the temporal evolution of the profiles during the AWH simulations and illustrates convergence.

In contrast, the position-dependent diffusion coefficient of ethanol was similar in the two membrane models (Fig. 6b). In both ACD and SCD, ethanol diffusion was reduced in the interfacial region between the lipid headgroups and the membrane core, whereas mobility increased within the hydrocarbon core. This enhanced mobility in the membrane interior is consistent with greater acyl-chain disorder and a higher density of three-dimensional packing defects in this region (see below). Thus, the lower ethanol permeability of ACD is primarily associated with differences in the free-energy barrier, rather than with reduced ethanol mobility along the permeation pathway.

### Hydrophobic defects

We next quantified membrane packing defects using complementary two- and three-dimensional analyses. Two-dimensional hydrophobic defects report water-exposed apolar lipid-tail atoms at the membrane surface [47], whereas three-dimensional packing defects identify internal voids that are free of atoms and large enough to accommodate a water-sized probe with a radius of 1.4 Å [48]. Thus, 2D defects quantify exposed hydrophobic surface, whereas 3D defects capture internal free volume independent of direct water exposure. Methodological details are provided in Materials and Methods, *Hydrophobic Defects*.

For deep hydrophobic defects, both asymmetric huRBC membrane models showed smaller defects in the exoplasmic leaflet than in the corresponding cytoplasmic leaflet (Fig. 7a). This leaflet asymmetry is also evident in representative ACD snapshots, in which deep defects are more abundant in the cytoplasmic leaflet than in the exoplasmic leaflet (Fig. 7c,d, red spheres). Thus, the more ordered and densely packed exoplasmic leaflets suppress deep exposure of apolar lipid-tail regions.

**Fig. 7.**
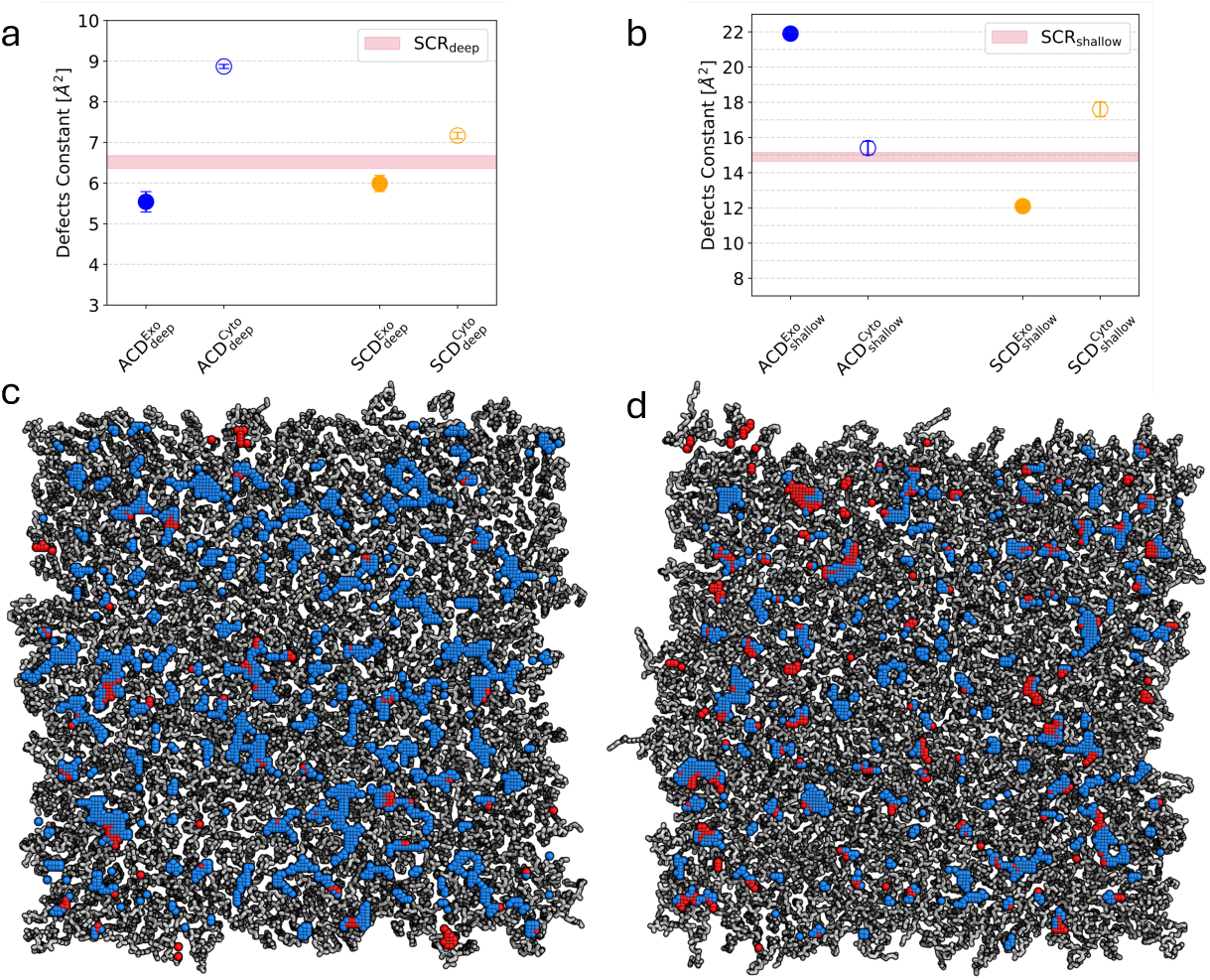
Two-dimensional membrane hydrophobic defects. (a,b) Characteristic defect size scale for deep defects (a), defined as defects extending more than 1 Å below the central phospholipid glycerol atom, and shallow defects (b), defined as defects located less than 1 Å below this reference position. Neighboring defect points were clustered into individual defects, and the resulting defect-area distribution, *p*(*A*), was fitted with an exponential decay, *p*(*A*) ∝ exp(*−A/*Π), where Π is the packing-defect constant. Circles denote ACD (blue) and SCD (orange), and the crimson line denotes SCR. (c,d) Representative snapshots of the ACD exoplasmic (c) and cytoplasmic (d) leaflets. Shallow and deep defects are shown as marine and red spheres, respectively.

Shallow defects showed a different behavior. In ACD, the exoplasmic leaflet exhibited larger shallow defects than the cytoplasmic leaflet, with a defect constant of approximately 22 Å^2^ compared with approximately 15.5 Å^2^ (Fig. 7b). The total shallow exposed hydrophobic surface was also larger in the ACD exoplasmic leaflet, amounting to approximately 2343 Å^2^, or 8.12% of the membrane surface, compared with approximately 1444 Å^2^, or 4.95%, in the cytoplasmic leaflet. This enhanced shallow hydrophobic exposure likely reflects the high cholesterol content of the ACD exoplasmic leaflet. Because cholesterol exposes only a small polar hydroxyl group at the membrane interface, it provides less headgroup shielding than phospholipids and can thereby increase hydrophobic exposure near the membrane surface.

By contrast, the SCD model exposed substantially less apolar surface within the exoplasmic leaflet. The SCD exoplasmic leaflet had a shallow-defect constant of approximately 12 Å^2^ and an exposed hydrophobic surface area of approximately 1191 Å^2^, corresponding to 4.53% of the membrane surface. The SCD cytoplasmic leaflet showed a larger shallow-defect area of approximately 1590 Å^2^, or 6.13%. In SCR, shallow hydrophobic exposure was nearly symmetric between leaflets (approximately 5.43% and 5.40% of the membrane surface exposed, respectively). These results indicate that shallow surface defects are strongly influenced by leaflet cholesterol content and interfacial shielding, whereas deep defects more closely follow leaflet packing and acyl-chain order.

A 3D packing-defect analysis further highlighted the strong leaflet imbalance in ACD (Fig. 8). Among the exoplasmic leaflets, ACD showed the lowest density of 3D packing defects, consistent with its small APL, high acyl-chain order, and slow lipid diffusion. Conversely, the ACD cytoplasmic leaflet showed the highest density of 3D defects, consistent with its larger APL, lower acyl-chain order, and faster lipid diffusion. Both asymmetric models showed leaflet-dependent differences in 3D packing defects, but the effect was strongest in ACD.

**Fig. 8.**
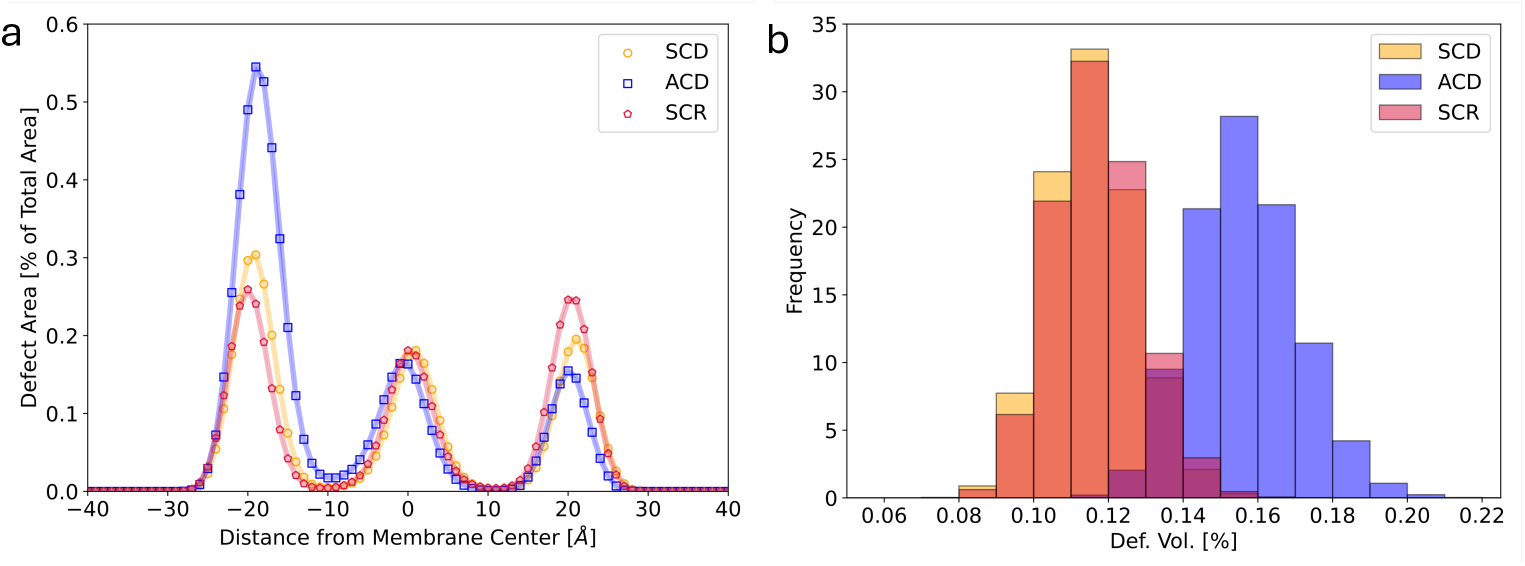
Three-dimensional membrane packing defects. (a) Distribution of total packing-defect area as a function of distance from the bilayer center of mass. Positive and negative distances correspond to the exoplasmic and cytoplasmic sides, respectively. (b) Distribution of total packing-defect volume for each membrane model. Data are normalized by the total membrane area in panel (a) and by the total membrane volume in panel (b).

The packing-defect volume distribution (Fig. 8b) correlated with the average water content in the corresponding leaflet regions (Fig. S2b). Larger defect volumes were associated with increased water occupancy, supporting the interpretation that internal packing defects facilitate water penetration into the membrane interior. Among the three models, ACD showed the largest defect-volume fraction relative to the total membrane volume. Together, the 2D and 3D defect analyses show that ACD combines reduced deep-defect formation in its densely packed exoplasmic leaflet with enhanced shallow hydrophobic exposure at the exoplasmic surface and increased internal free volume on the cytoplasmic side.

### Cholesterol clustering

To assess whether cholesterol organization contributes to hydrophobic defect formation, we quantified leaflet-dependent cholesterol clustering in each membrane model. The clustering analysis is described in Materials and Methods, *Cholesterol Clustering*.

In most systems and leaflets, less than 30% of cholesterol molecules were assigned to clusters (Fig. 9a). The clear exception was the ACD exoplasmic leaflet, where approximately 68% of cholesterol molecules were clustered. This pronounced clustering is consistent with the high cholesterol-to-phospholipid ratio in this leaflet, which exceeds 1:1 in favor of cholesterol (Table 3). The high local cholesterol density also led to transient formation of very large clusters, in some cases containing one third or more of all cholesterol molecules in the leaflet (Fig. 9b,c). Although such large clusters are rare, as indicated by the logarithmic cluster-size distribution, they reveal transient regions of very high cholesterol density. No crystalline cholesterol ordering was observed in the present simulations. Overall, clustered cholesterol occupied approximately 37% of the ACD exoplasmic leaflet area (17% by free cholesterol, only 46% by phospholipids; Fig. 10a).

**Fig. 9.**
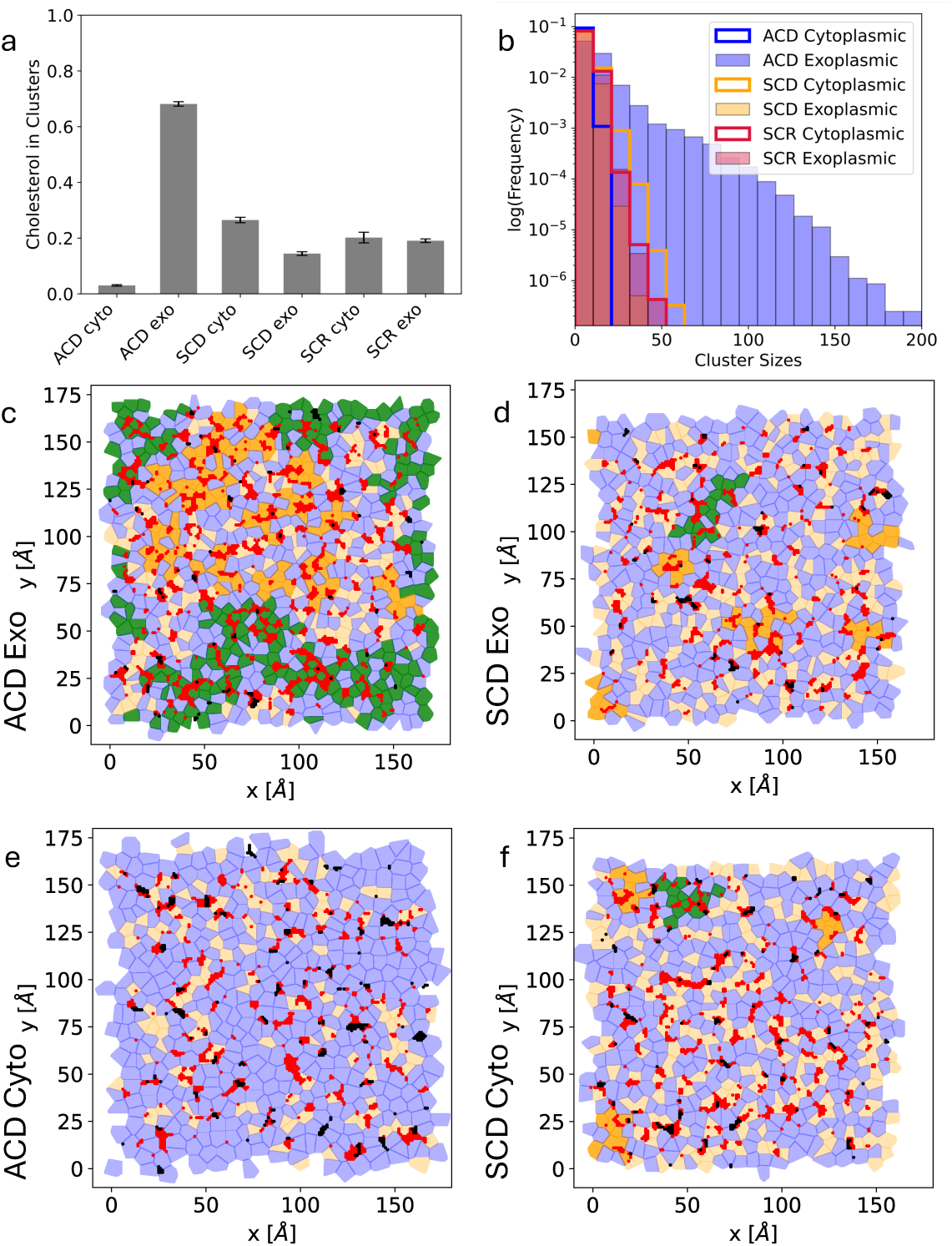
Cholesterol clustering. (a) Average fraction of cholesterol molecules assigned to clusters. (b) Distribution of cholesterol cluster sizes for each membrane model and leaflet, shown on a logarithmic scale. (c,d,e,f) Voronoi representations of the ACD exoplasmic (c), ACD cytoplasmic (e), SCD exoplasmic (d) and SCD cytoplasmic (f) leaflets for a representative frame. Phospholipids are shown in blue, cholesterol molecules belonging to smaller clusters in dark orange, and non-clustered cholesterol molecules in very light orange. The largest cholesterol clusters are shown in dark green. In panel (c), the largest cholesterol cluster contains approximately 180 molecules. Red and black dots indicate shallow and deep 2D hydrophobic defects, respectively.

**Fig. 10.**
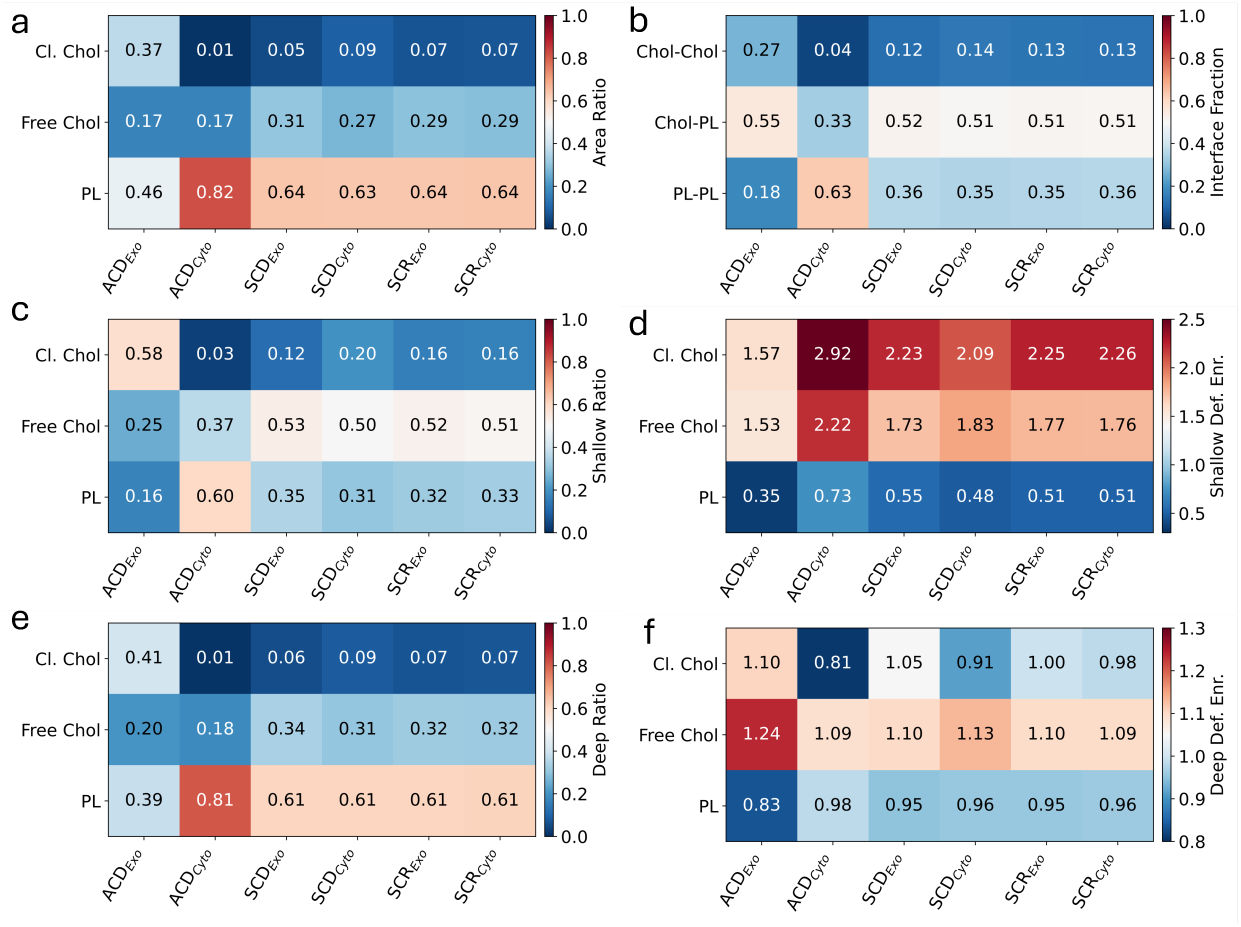
Enrichment of 2D hydrophobic defects in cholesterol- and phospholipid-occupied regions. (a) Fraction of the leaflet area occupied by clustered cholesterol (Cl. Chol), nonclustered cholesterol (Free Chol), and phospholipids (PL). (b) Fraction of 2D Voronoi nearest-neighbor interfaces assigned to cholesterol–cholesterol (Chol—Chol), cholesterol–phospholipid (Chol—PL), and phospholipid–phospholipid (PL—PL) contacts. (c) Fraction of shallow defects located within each region, relative to the total number of shallow defects in the respective leaflet. (d) Enrichment indices for shallow defects in Voronoi regions occupied by Cl.Chol, Free Chol, and PL. Values above one indicate enrichment, whereas values below one indicate depletion. (e) Fraction of deep defects located within each region, relative to the total number of deep defects in the respective leaflet. (f) Corresponding enrichment indices for deep defects.

The Voronoi representations in Fig. 9c-f suggest a spatial association between cholesterol-rich regions and 2D hydrophobic defects. We quantified this association using the defect enrichment index (Fig. 10d,f), 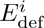 (see Materials and Methods, *Cholesterol Clustering*). This metric reports whether defects occur preferentially in Voronoi regions occupied by clustered cholesterol, non-clustered cholesterol, or phospholipids. Because 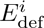 is normalized by the area occupied by each lipid category, it is best suited for comparing defect densities among lipid environments within the same leaflet. It should not be interpreted as a direct measure of absolute defect abundance across different leaflets.

Shallow hydrophobic defects were consistently enriched in cholesterol-occupied regions and depleted in phospholipid regions across all membrane models and leaflets (Fig. 10d). This enrichment was observed for both non-clustered cholesterol and clustered cholesterol, and was strongest in clustered cholesterol regions. Thus, cholesterol-rich regions promote shallow hydrophobic exposure at the membrane surface, consistent with the limited polar shielding provided by the small cholesterol hydroxyl group.

In turn, deep defects showed weaker and less systematic associations with cholesterol organization (Fig. 10f). In most cases, deep defects were only modestly enriched near non-clustered cholesterol and showed no comparably strong enrichment in clustered cholesterol regions. These data indicate that cholesterol clustering primarily affects shallow surface defects, whereas deep hydrophobic defects are more strongly governed by leaflet packing, acyl-chain order, and internal free volume.

The ACD exoplasmic leaflet, which showed the largest shallow-defect constant (Fig. 7b), also had the highest fraction of cholesterol–cholesterol interfaces, approximately 27% (Fig. 10b), while the fraction of cholesterol–phospholipid interfaces was only slightly increased compared to SCD or SCR models. This reflects saturation of cholesterol–phospholipid mixing and accounts for the enrichment of shallow defects in cholesterol-rich regions. Conversely, the ACD cytoplasmic leaflet showed the largest number of deep defects and the largest deep-defect constant, consistent with its loose packing, low acyl-chain order, and low cholesterol content. Thus, cholesterol clustering primarily promotes shallow hydrophobic exposure, whereas deep defects are controlled mainly by leaflet packing and order.

### Local lipid environment

We next quantified the local lipid environment of each lipid species using an enrichment index that compares the observed nearest-neighbor composition with that expected for random mixing (see Materials and Methods, *Local Lipid Environment*). This analysis identifies lipid-specific neighborhood preferences within each leaflet and complements the cholesterol-clustering analysis described above.

Across the PM models, cholesterol showed only limited pairwise self-enrichment in the nearest-neighbor analysis, indicating that cholesterol generally contacts phospholipids more frequently than other cholesterol molecules (Fig. 11; Fig. S3). This observation does not contradict the cholesterol-clustering analysis, because the enrichment matrices report local composition relative to each leaflet composition, whereas clustering identifies contiguous cholesterol-rich regions based on spatial connectivity. In particular, the ACD exoplasmic leaflet contains large cholesterol clusters (Fig. 9), but its nearest-neighbor enrichment matrix does not indicate strong selectivity for a specific phospholipid partner.

**Fig. 11.**
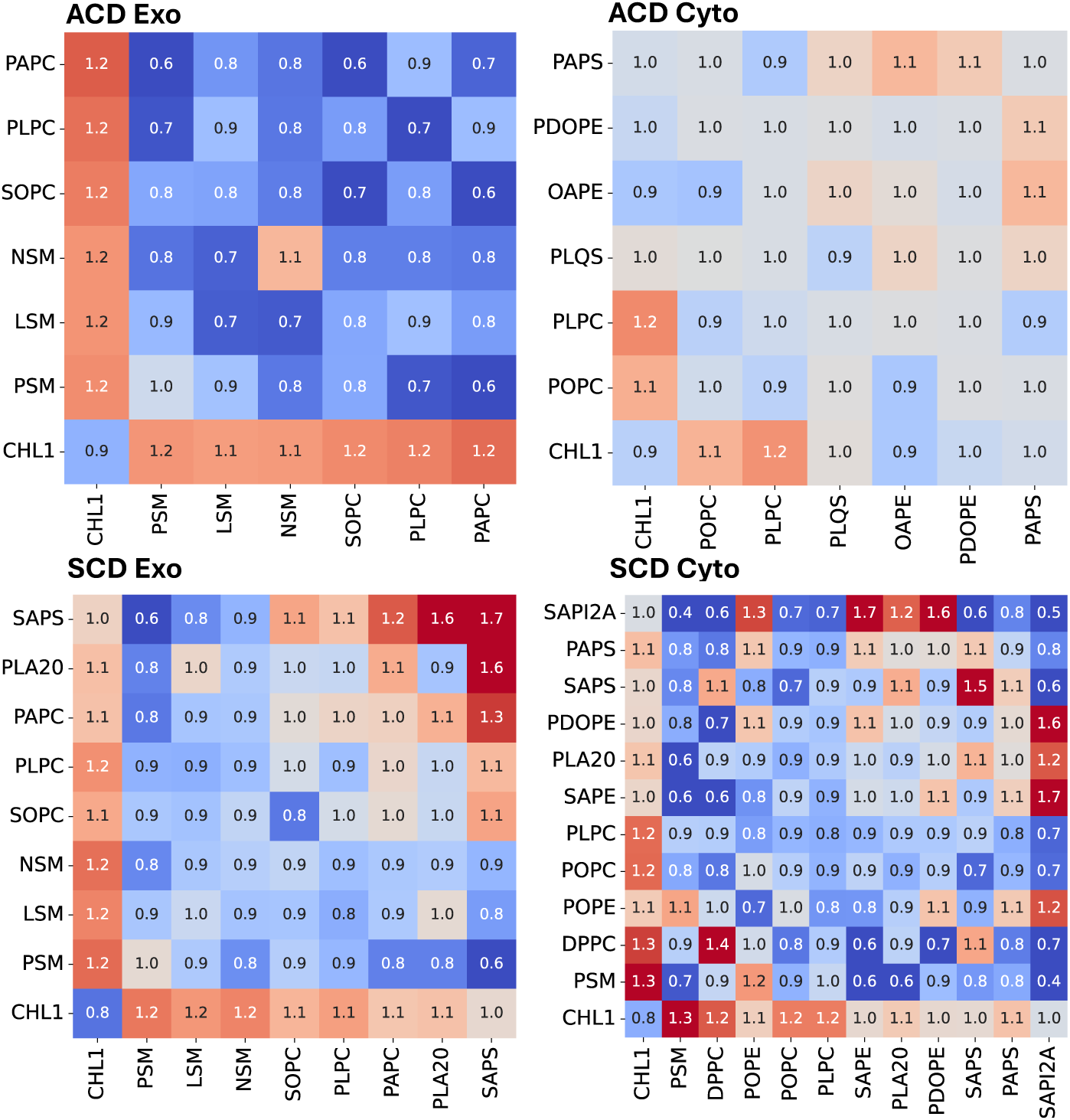
Lipid enrichment matrices for the ACD and SCD membrane models. Enrichment indices were computed for each lipid species within each leaflet. The *y*-axis indicates the reference lipid type, and the *x*-axis indicates the neighboring lipid type. Values greater than one indicate enrichment relative to a random mixture, whereas values below one indicate depletion. Shown are the exoplasmic (Exo) and cytoplasmic (Cyto) leaflets of the ACD and SCD models. Lipids are ordered by degree of unsaturation and charge.

In both huRBC membrane models, cholesterol showed local association with SM- and saturated-lipid-rich environments (Fig. 11). Consistently, the cholesterol–PSM radial distribution function (RDF) displayed a more compact first solvation shell than the cholesterol–PLPC RDF (Fig. S4). This result indicates tighter cholesterol–SM packing than cholesterol–PC packing, in line with favorable sterol–sphingomyelin interactions. However, neither the enrichment matrices nor the RDFs provide evidence for stable SM/saturated-lipid–cholesterol domain formation beyond local molecular-scale contacts.

The ACD exoplasmic leaflet showed a distinct behavior because of its high cholesterol content. In this leaflet, cholesterol contacts were frequent even with unsaturated phospholipids, despite their less favorable packing complementarity with cholesterol. This behavior reflects the high cholesterol-to-phospholipid ratio in the ACD exoplasmic leaflet, which increases the probability of cholesterol contacts with all available phospholipid classes, including the relatively small pool of unsaturated species.

In contrast, the ACD cytoplasmic leaflet showed nearly random lipid mixing. Its enrichment matrix was comparatively featureless, consistent with its high fluidity, large APL, low acyl-chain order, and rapid lipid diffusion. The SCD cytoplasmic leaflet, and similarly the SCR model (Fig. S3), displayed more structured enrichment matrices, indicating stronger local lipid-neighborhood heterogeneity. Negatively charged lipids, including phosphatidylserine and phosphatidylinositol species such as SAPI2A, contributed prominently to this heterogeneity together with neutral polyunsaturated lipids.

Despite its low abundance, SAPI2A (i.e. PIP_2_) formed a distinct local environment in the cytoplasmic leaflet, likely driven by van der Waals interactions between lipid tails. This finding suggests that minor signaling lipids can generate specific local neighborhoods even in compositionally complex membranes. Such lipids should therefore be retained in realistic plasma membrane models when local membrane organization, protein recruitment, or signaling-relevant lipid environments are investigated. Overall, the enrichment analysis shows that leaflet composition and lipid unsaturation shape local lipid neighborhoods, whereas cholesterol-rich regions primarily affect local packing and hydrophobic exposure rather than producing large, compositionally selective domains.

### Area compressibility and bending

We next quantified the elastic properties of the membrane models. The area compressibility modulus, *K_a_*, was obtained from box-area fluctuations (BAF; see Materials and Methods, *Area Compressibility*) and compared with experimental and simulation values reported for simpler synthetic membranes [49–52] (Fig. 12).

**Fig. 12.**
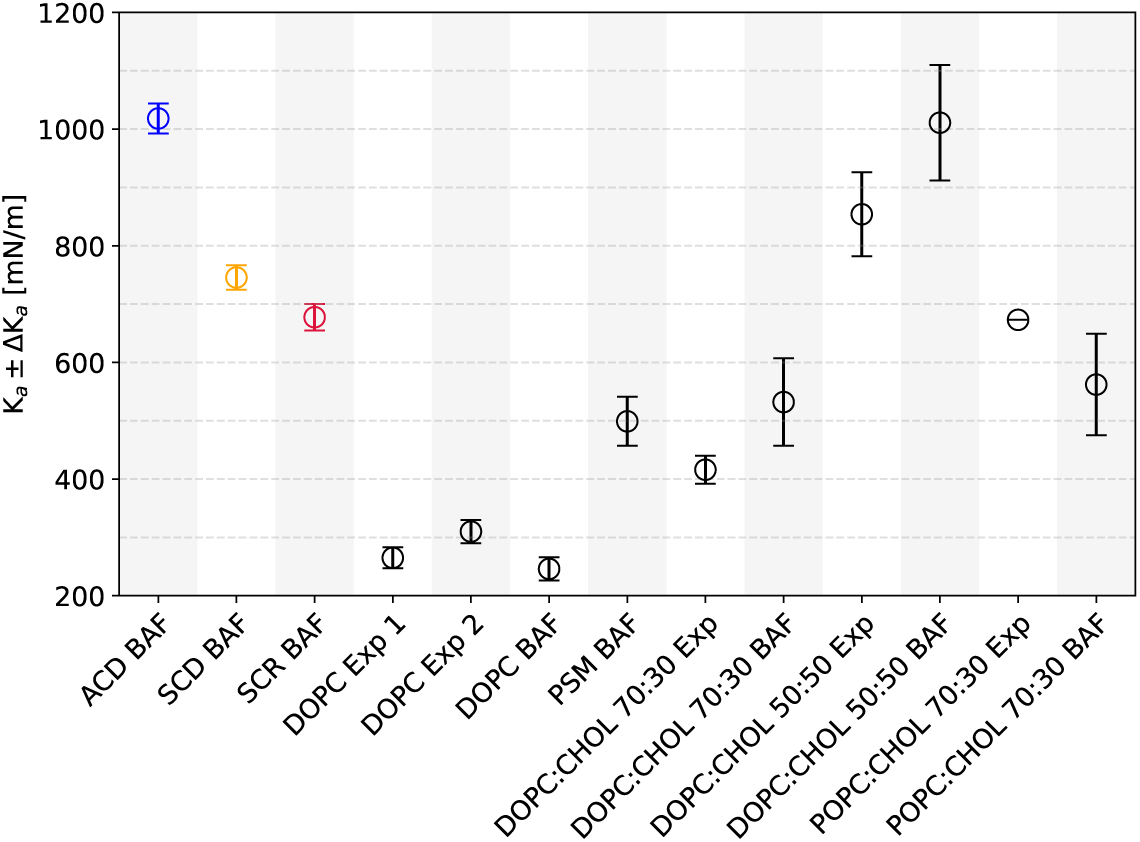
Area compressibility moduli. Area compressibility moduli, *Ka*, for the ACD (blue), SCD (orange), and SCR (crimson) plasma membrane models, shown together with literature values for synthetic membranes (black). Error bars for the present simulations were estimated by block analysis of the box area followed by error propagation. The x-axis lists the membrane models and indicates whether *Ka* was obtained experimentally (Exp) or from simulation box area fluctuations (BAF). Data origin: ACD BAF, SCD BAF, SCR BAF - own computation; DOPC Exp 1 - Rawicz *et al.*, 2000 [50]; DOPC Exp 2 - Rawicz *et al.*, 2008 [51]; PSM BAF, DOPC:CHOL 70:30 BAF, DOPC:CHOL 50:50 BAF, POPC:CHOL 70:30 - Doktorova *et al.*, 2019 [49]; DOPC:CHOL 70:30 Exp, DOPC:CHOL 50:50 Exp, POPC:CHOL 70:30 Exp - Evans *et al.*, 2013 [52].

Among the three plasma membrane models, ACD had the highest area compressibility modulus and was therefore the least laterally compressible (Fig. 12). SCD was also less compressible than SCR, but the difference between SCD and SCR was small compared with the increase observed for ACD. The *K_a_* value of ACD was comparable to values reported for cholesterol-rich symmetric bilayers with cholesterol-to-phospholipid ratios similar to that of the ACD exoplasmic leaflet (PL:CHOL 44:56 and DOPC:CHOL 50:50).

The high *K_a_* of ACD is consistent with the dense packing of its exoplasmic leaflet (Fig. 2). This leaflet has the smallest APL among all leaflets analyzed and is therefore expected to suppress lateral area fluctuations, as further compression becomes sterically unfavorable. The measured *K_a_* of ACD is likely dominated by the densely packed, weakly compressible exoplasmic leaflet, whereas the higher compressibility of the more fluid cytoplasmic leaflet is not resolved separately.

We then quantified bending rigidity using the bending modulus, *κ_b_*, obtained from the local fluctuation method (LFM [32]; see Materials and Methods, *Bending*; Fig. 13). The bending moduli followed the same ordering as 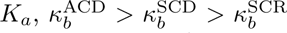. However, the differences were small. ACD showed the highest value, *κ_b_* = (38.1 ± 0.1) *k_B_T*, whereas SCD and SCR yielded *κ_b_* = (36.6±0.1) *k_B_T* and *κ_b_* = (36.2±0.1) *k_B_T*, respectively (Fig. 13). Thus, strong phospholipid-number asymmetry substantially increases resistance to lateral compression but only modestly increases the apparent bending rigidity.

**Fig. 13.**
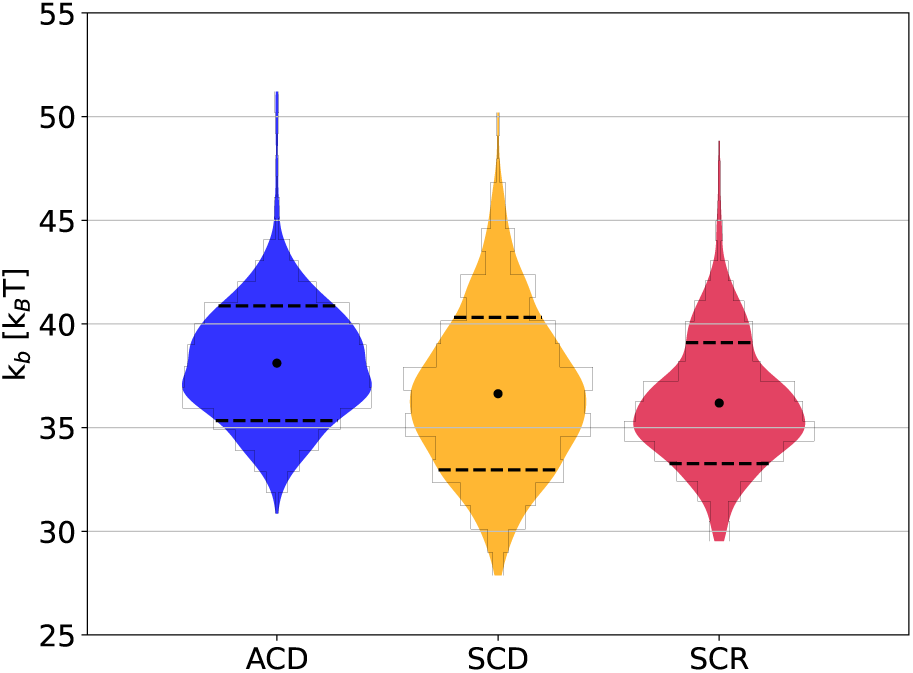
Bending moduli obtained using the local fluctuation method (LFM). Bending moduli, *κ_b_*, are reported in units of *k_B_T* for the ACD (38.1 *±* 0.1 *k_B_T*), SCD (36.6 *±* 0.1 *k_B_T*), and SCR (36.2 *±* 0.1 *k_B_T*) models (x-axis), calculated using circular patches of radius *r* = 3.0 nm (see Materials and Methods, *Bending*). Black dots indicate the mean of the raw bending-modulus distributions, shown as open histogram bars, and dashed lines indicate the standard deviation. Filled violins show Gaussian kernel density estimates of the raw distributions for the ACD (blue), SCD (orange), and SCR (crimson) models. Distributions were obtained from 1000 independent LFM evaluations using randomly selected patch centers.

This weaker effect on *κ_b_* reflects the different mechanical modes probed by area compression and bending. Area compression is particularly sensitive to the least compressible leaflet, whereas bending requires coordinated deformation of both leaflets. In ACD, the densely packed exoplasmic leaflet increases resistance to deformation, but the more fluid cytoplasmic leaflet can partly accommodate curvature and local height fluctuations. Consequently, the pronounced leaflet-packing asymmetry of ACD strongly suppresses projected-area fluctuations but produces only a moderate increase in bending rigidity.

## Discussion

Plasma membranes combine pronounced lipid-type asymmetry with an unresolved degree of phospholipid-number asymmetry between the two leaflets. Here, we used atomistic molecular dynamics simulations to compare two human red blood cell (huRBC) plasma membrane models that differ primarily in this latter property. The ACD model was based on a recent lipidomics study by Doktorova *et al.*, which proposed a 50% overabundance of phospholipids in the cytoplasmic leaflet and compensatory cholesterol enrichment in the exoplasmic leaflet [10]. The SCD model, by contrast, retains strong lipid-type asymmetry but contains similar phospholipid and cholesterol numbers in the two leaflets [9]. A scrambled membrane served as a symmetric reference. This comparison allowed us to separate the effects of lipid-type asymmetry from those caused by strong phospholipid-number asymmetry.

Both asymmetric models reproduced the canonical organization of the plasma membrane: a more ordered and more densely packed exoplasmic leaflet, and a more disordered and more fluid cytoplasmic leaflet. However, the magnitude of this transbilayer contrast differed strongly between models. The ACD model (with 30% phospholipid overabundance in cytoplasmic leaflet) generated an exceptionally compressed, cholesterol-rich exoplasmic leaflet and a comparatively dilute cytoplasmic leaflet. This resulted in large cholesterol-rich clusters, strongly reduced exoplasmic lipid mobility, altered hydrophobic-defect distributions, reduced water and ethanol permeability, and a substantially increased area compressibility modulus. By contrast, the SCD model preserved the same qualitative leaflet asymmetry but avoided several extreme features of ACD, including large cholesterol domains and the order-of-magnitude interleaflet diffusion imbalance.

The simulations do not identify a single model as universally correct. Rather, they define the physical consequences of imposing strong phospholipid-number asymmetry. Some observables, especially lipid diffusion and ethanol permeability, are more closely reproduced by the SCD model. Our comparison suggests that key features of plasma membrane asymmetry arise from lipid-type asymmetry alone, without requiring a large phospholipid-number imbalance. Moderate cholesterol asymmetry or moderate leaflet-density differences may nevertheless remain important features of biological plasma membranes.

### Experimental constraints do not uniquely define phospholipid-number asymmetry

The experimental basis for a large phospholipid-number imbalance remains uncertain. However, previous estimates of phospholipid transbilayer distributions in huRBCs based on chemical or enzymatic accessibility assays consistently support lipid-type asymmetry, with sphingomyelin and phosphatidylcholine enriched in the exoplasmic leaflet and phosphatidylethanolamine (PE), phosphatidylserine (PS), phosphatidylinositol (PI), and phosphorylated phosphoinositides enriched in the cytoplasmic leaflet [4–10]. Quantitatively, however, reported leaflet distributions differ substantially (Table S4). This uncertainty is critical because PC and PE are abundant lipid classes; even moderate changes in their inferred exoplasmic fractions strongly affect the calculated phospholipid numbers in the two leaflets.

Several methodological factors may contribute to this variability. Phospholipase A_2_ (PLA_2_)-based assays rely on efficient hydrolysis of exoplasmic phospholipids containing an ester-linked fatty acid at the *sn*-2 position. However, incomplete digestion of exoplasmic PC can lead to an overestimation of the cytoplasmic PC fraction [33]. PLA_2_ activity may also be affected by product inhibition, enzyme source, and the physical state of the membrane [33, 34, 53]. In particular, PLA_2_ from *Apis mellifera* has been reported to digest PC inefficiently in huRBC membranes [33, 35]. Efficient digestion may require reduced exoplasmic leaflet tension, for example by sequential sphingomyelinase treatment [35]. These issues have motivated recent criticism of strongly asymmetric phospholipid number models [33, 34].

Here, we combined representative total huRBC lipid compositions with selected leafletdistribution estimates (Table 2). The latter are based on the studies of Verkleij *et al.* and van Meer *et al.*, which employed *Naja naja* PLA_2_ rather than bee venom-derived PLA_2_ and included conditions in which sphingomyelinase treatment reduces leaflet tension [5, 7, 35]. These studies also report a finite exoplasmic PE fraction, consistent with the independent TNBS-based estimate of Gordesky and Marinetti [54]. Approximate distributions of 80% SM, 80% PC, 20% PE, and 0% PS/PI/PA in the exoplasmic leaflet were inferred from these studies. Using these lipid fractions, the phospholipid-number ratio between cytoplasmic and exoplasmic leaflets is close to unity for several classical huRBC compositions: *R*_PL_ ≈ 1.05 for Verkleij *et al.* [5] and *R*_PL_ ≈ 1.04 for van Meer *et al.* [7]. Applying the same leaflet fractions to the Lorent *et al.* lipidome gives a stronger cytoplasmic enrichment, *R*_PL_ ≈ 1.31 [9], whereas the Doktorova *et al.* composition gives *R*_PL_ ≈ 1.11 under this conservative assignment [10].

**Table 1.** Approximate phospholipid distributions across the two leaflets for representative human red blood cell lipid compositions. Values are given in mol% of total phospholipids. The exoplasmic and cytoplasmic rows were estimated using approximate leaflet distributions of 80% SM, 80% PC, 20% PE/PEp, and 0% PS/PI/PA in the exoplasmic leaflet. Glycosphingolipids (GSLs) were explicitly included only where reported. PEp denotes plasmalogen phosphatidylethanolamine. PS, PI, and phosphatidic acid (PA) are grouped as minor anionic lipids. Cholesterol is not included in the phospholipid totals.

| Reference | Leaflet | SM | PC | GSL | PE/PEp | PS/PI/PA | Total |
| --- | --- | --- | --- | --- | --- | --- | --- |
| Verkleij *et al.* [5] | Both | 23.5 | 29.4 | – | 31.9 | 15.2 | 100.0 |
|  | Exo | 18.8 | 23.5 | – | 6.4 | 0.0 | 48.7 |
|  | Cyto | 4.7 | 5.9 | – | 25.5 | 15.2 | 51.3 |
| van Meer *et al.* [7] | Both | 25.3 | 29.5 | – | 25.9 | 19.3 | 100.0 |
|  | Exo | 20.2 | 23.6 | – | 5.2 | 0.0 | 49.0 |
|  | Cyto | 5.1 | 5.9 | – | 20.7 | 19.3 | 51.0 |
| Lorent *et al.* [9] | Both | 16.8 | 30.3 | – | 27.6 | 25.3 | 100.0 |
|  | Exo | 13.5 | 24.2 | – | 5.5 | 0.0 | 43.2 |
|  | Cyto | 3.3 | 6.1 | – | 22.1 | 25.3 | 56.8 |
| Doktorova *et al.* [10] | Both | 16.2 | 30.3 | 4.7 | 27.2 | 21.6 | 100.0 |
|  | Exo | 13.0 | 24.2 | 4.7 | 5.4 | 0.0 | 47.3 |
|  | Cyto | 3.2 | 6.1 | 0.0 | 21.8 | 21.6 | 52.7 |

**Table 2.** Diffusion coefficient ratios used to compare experimental and simulated effects of plasma membrane asymmetry. Experimental values report the ratio between asymmetric and scrambled fibroblast plasma membranes measured for TopFluor-sphingomyelin (SM) and TopFluor-cholesterol (Chol) [10]. Simulation values report the corresponding ratios between the exoplasmic leaflet of the asymmetric plasma membrane models, ACDexo or SCDexo, and the scrambled reference membrane, SCR.

| Method | Lipid | Comparison | $D_A/D_B$ |
| --- | --- | --- | --- |
| Experiment [10] | SM | Asymmetric / scrambled fibroblast PM | $0.60 \pm 0.10$ |
| | Chol | Asymmetric / scrambled fibroblast PM | $0.82 \pm 0.22$ |
| AA-MD | SM | ACD<sub>exo</sub> / SCR | $0.26 \pm 0.01$ |
| | Chol | ACD<sub>exo</sub> / SCR | $0.24 \pm 0.01$ |
| | SM | SCD<sub>exo</sub> / SCR | $0.63 \pm 0.02$ |
| | Chol | SCD<sub>exo</sub> / SCR | $0.60 \pm 0.02$ |

The available lipidomic compositions are compatible with near-symmetric to moderately asymmetric phospholipid numbers, depending on the assumed PC and PE leaflet distributions. In summary, the experimental literature robustly supports lipid-type asymmetry but does not yet define a unique phospholipid-number asymmetry. This distinction is central for interpreting the present simulations. The ACD model represents a strong number-asymmetry scenario in which cholesterol redistribution compensates leaflet stress. The SCD model represents an alternative scenario in which lipid-type asymmetry is retained without a large phospholipid-number imbalance. Their comparison isolates which membrane properties follow specifically from strong phospholipid-number asymmetry rather than from lipid-type asymmetry alone.

### High exoplasmic cholesterol density drives cholesterol clustering and surface exposure

A central consequence of the ACD model is the high cholesterol content of the exoplasmic leaflet. Some cholesterol enrichment in the exoplasmic leaflet is plausible, because cholesterol preferentially associates with saturated lipids and sphingomyelin and can redistribute to compensate leaflet stress [13, 32, 55–59]. However, the ACD model drives this mechanism to an extreme, with an exoplasmic cholesterol fraction of approximately 56 mol%. At this concentration, cholesterol no longer behaves only as a condensing component dispersed among phospholipids [41]. Instead, the ACD exoplasmic leaflet forms large cholesterol-rich clusters. Although no crystalline cholesterol ordering was observed, the local cholesterol density approaches reported cholesterol miscibility limits in simplified phospholipid–cholesterol bilayers [60–62]. This suggests that even stronger phospholipid-number asymmetries would require cholesterol concentrations that may approach the boundary of bilayer miscibility and potentially favor cholesterol-rich or crystalline phases.

Cholesterol enrichment also reshaped the hydrophobic-defect landscape. In ACD, the densely packed exoplasmic leaflet was depleted in deep defects but enriched in shallow defects, and vice versa for the cytosolic leaflet. This distinction is important because different membrane-binding motifs sense different defect depths. ALPS motifs and several amphipathic peptides preferentially recognize deep defects that can accommodate bulky hydrophobic residues [63–67]. By contrast, smaller amphipathic motifs and some polar amphipathic polymers may be more sensitive to shallower defects [47, 68–70]. Thus, cholesterol enrichment in the ACD exoplasmic leaflet does not simply make the membrane uniformly less permissive to insertion. Instead, it shifts the defect spectrum toward shallow hydrophobic exposure at the extracellular side.

Despite the high abundance of cholesterol and sphingomyelin in the ACD exoplasmic leaflet, the simulations did not reveal large, compositionally selective sphingomyelin–cholesterol domains. Cholesterol–sphingomyelin contacts were geometrically tighter than cholesterol–phosphatidylcholine contacts, consistent with favorable sterol–sphingomyelin interactions, but the nearest-neighbor enrichment analysis did not support stable domain formation beyond local molecular-scale contacts. This apparent discrepancy likely reflects saturation of the local lipid environment: when cholesterol is highly abundant, cholesterol contacts become unavoidable, reducing the contrast between preferred and non-preferred neighbors. The strongest composition-dependent heterogeneity instead appeared in the SCD cytoplasmic leaflet and in the scrambled membrane, where polyunsaturated and anionic lipids, including phosphoinositide-containing species, formed distinct local environments. Thus, plasma membrane heterogeneity is not governed only by cholesterol and sphingomyelin but also by polyunsaturated and charged lipids. Conclusions on domain formation should, however, be interpreted with caution. The present models lack transmembrane and peripheral proteins, cortical actin, and membrane-associated scaffolds, all of which can organize or stabilize nanoscale membrane domains in cells [71–75]. Further experiments using probes with calibrated shallow-versus deep-defect preferences, or direct simulations of lipidated peptides in both models, would help resolve this ambiguity.

### Area per lipid, diffusion and permeability provide complementary experimental tests

Experimental observables support different aspects of the two models. The ACD exoplasmic leaflet gives the closest qualitative agreement with area-per-lipid values inferred from Di-4-ANEPPDHQ measurements [10]. However, this comparison is indirect. Di-4-ANEPPDHQ reports an environment-sensitive fluorescence response that correlates with lipid order and packing under calibrated conditions but is not determined by area per lipid alone [76, 77]. Agreement with inferred APL values should therefore not be treated as definitive validation of the ACD model.

Lipid diffusion provides a more sensitive dynamic readout of leaflet packing. In the simulations, ACD represents the most extreme case: lipids in the cholesterol-rich exoplasmic leaflet diffuse much more slowly than those in the cytoplasmic leaflet. This behavior follows from the small APL, high acyl-chain order, and high cholesterol density of the ACD exoplasmic leaflet. SCD shows the same qualitative direction of asymmetry but a more moderate interleaflet contrast, because it retains lipid-type asymmetry without imposing a large phospholipid-number imbalance.

Absolute diffusion coefficients are difficult to compare directly with experiment. Experimental measurements often use fluorescent lipid analogues, such as TopFluor-sphingomyelin and TopFluor-cholesterol, whereas the simulations report unmodified lipid species. Fluorescent labels can alter packing, partitioning, and interactions with neighboring lipids. In addition, all-atom simulations probe nanometer-scale motion on nanosecond to microsecond timescales, whereas FCS-based approaches report effective diffusion over larger spatiotemporal scales. Relative changes upon scrambling are therefore more informative than absolute diffusion coefficients.

In experiments, scrambling increased the diffusion of TopFluor-sphingomyelin and TopFluor-cholesterol, giving asymmetric/scrambled diffusion ratios of 0.60 ± 0.10 for SM and 0.82 ± 0.22 for cholesterol [10]. The corresponding simulation ratios for ACD_exo_/SCR were much smaller, 0.26 ± 0.01 for SM and 0.24 ± 0.01 for cholesterol, indicating that ACD predicts a stronger exoplasmic mobility suppression than observed experimentally. By contrast, the SCD_exo_/SCR ratios, 0.63 ± 0.02 for SM and 0.60±0.02 for cholesterol, are closer to the experimental scrambling response, although cholesterol diffusion remains underestimated relative to the experimental ratio.

Leaflet-resolved diffusion leads to the same conclusion. ACD predicts approximately 12-fold faster phospholipid diffusion and approximately 14-fold faster cholesterol diffusion in the cytoplasmic leaflet than in the exoplasmic leaflet. Such a large interleaflet mobility difference is difficult to reconcile with available leaflet-resolved measurements. Gupta *et al.*, for example, reported faster diffusion in the cytoplasmic leaflet than in the exoplasmic leaflet, but the difference was closer to a factor of two for TopFluor-PC in CHO-K1 and RBL-2H3 cells [78]. The SCD model, with an interleaflet diffusion ratio of approximately 2.5–3, is close to this experimental scale. Thus, diffusion supports a more fluid cytoplasmic leaflet but argues against the very large dynamical asymmetry produced by the ACD composition tested here.

A similar picture is provided by membrane permeability: Ethanol permeability favors the SCD model. Brahm measured an ethanol permeability coefficient of *K_p_* = 2.1 × 10*^−^*^3^ cm/s for huRBCs at 298 K [79]. The SCD model gives *K_p_* = (1.70 ± 0.05) × 10*^−^*^3^ cm/s at 310 K, whereas the ACD model gives a much lower value of *K_p_*= (0.23 ± 0.05) × 10*^−^*^3^ cm/s. Although the temperature difference limits direct quantitative comparison, higher temperature would generally be expected to increase, not decrease, small-solute permeation. The approximately seven- to eightfold lower ethanol permeability of ACD is therefore unlikely to be explained by temperature alone and instead reflects the high free-energy barrier associated with its densely packed exoplasmic leaflet. Noteworthy, a similarly reduced permeation across the ACD model compared to SCD was observed for water molecules.

Taken together, diffusion and permeability suggest that lipid-type asymmetry is sufficient to generate a mobility landscape in agreement with experiment, whereas strong phospholipid-number asymmetry likely over-amplifies exoplasmic packing and plasma membrane barrier properties.

### Mechanical properties distinguish area compression from bending

Mechanical properties further showed that cholesterol distribution, not only total cholesterol abundance, controls membrane elasticity. The ACD membrane exhibits the highest area compressibility modulus (*>* 1000 mN/m), corresponding to an approximately 40% lower lateral compressibility than SCD (Figure 12), indicating that the cholesterol-rich exoplasmic leaflet strongly suppresses area fluctuations. Because ACD, SCD, and SCR contain similar total cholesterol contents, this increase arises from cholesterol redistribution and leaflet packing rather than cholesterol abundance alone. In an asymmetric bilayer, box-area fluctuations inform about the response of the coupled membrane and are likely dominated by the least compressible leaflet. The more fluid ACD cytoplasmic leaflet may therefore be masked by the densely packed exoplasmic leaflet in the apparent *K_a_*. Direct experimental validation of *K_a_*is difficult. Micropipette aspiration measurements on osmotically preswollen red blood cells report apparent area compressibility moduli of 288 ± 50 mN/m at 25*^◦^*C and 350 mN/m at 40,*^◦^*C [80, 81]. These values, however, describe the composite response of the lipid bilayer, spectrin network, and available excess membrane area rather than the intrinsic lipid-bilayer compressibility alone. Similar considerations apply to vesicle measurements, where suppression of thermal undulations and recruitment of hidden area reservoirs can make the apparent modulus, 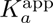, smaller than the intrinsic stretching modulus, 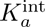 [50]. Thus, experimental area-compressibility values provide useful scale estimates but are not sufficient to discriminate unambiguously between the ACD and SCD models.

Bending moduli *κ_b_*followed the same ordering as the compressibility modulus,

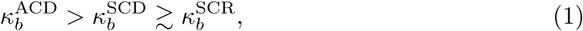

but the differences were much smaller (Figure 13). A close similarity between the bending modulus of a compositionally asymmetric model membrane and its symmetrized counterpart has also been reported in previous computational studies for simplified model systems [82, 83]. This weaker sensitivity is physically plausible because bending requires coordinated deformation of both leaflets, allowing the more fluid cytoplasmic leaflet in ACD to partly compensate for the rigid exoplasmic leaflet. However, similar to *K_a_*, experimental validation remains challenging [50, 80, 81]. Reported bending moduli measured using the same experimental technique (flickering analysis) range from approximately 3 *k*_B_*T* [84] to 210 *k*_B_*T* [85]. Thus, the bending moduli determined here are physically reasonable but do not allow discrimination between ACD and SCD (Table S3).

### Conclusions

Our simulations show that phospholipid-number asymmetry strongly amplifies structural and dynamical differences between the leaflets of huRBC plasma membranes. In the ACD model, cytoplasmic phospholipid excess drives cholesterol enrichment in the exoplasmic leaflet [10], leading to dense lipid packing, large cholesterol-rich clusters, increased extracellular exposed hydrophobic surface, reduced water and ethanol permeability, and increased lateral stiffness. These effects demonstrate that leaflet number-density imbalance is a powerful determinant of plasma membrane material properties.

However, several consequences of the ACD model appear more pronounced than suggested by available experimental constraints. In particular, ACD predicts an order-of-magnitude interleaflet diffusion imbalance and a substantially reduced ethanol permeability, whereas the SCD model more closely reproduces these dynamic observables while retaining the canonical plasma membrane organization of an ordered exoplasmic leaflet and a fluid cytoplasmic leaflet. Thus, lipid-type asymmetry alone can account for many hallmark features of plasma membrane asymmetry without requiring a large phospholipid-number imbalance.

We therefore conclude that a strongly asymmetric phospholipid number density is not required to reproduce key physical properties of the plasma membrane. Instead, biological plasma membranes may be better described by a moderate combination of lipid-type asymmetry, cholesterol asymmetry, and leaflet-density imbalance.

## Materials and Methods

### All-atom simulations

We performed all-atom (AA) molecular dynamics (MD) simulations of complex plasma membrane (PM)-mimetic bilayers with symmetric and asymmetric leaflet compositions. The bilayers were constructed using experimentally motivated lipid compositions and leaflet distributions (Table 3), solvated in 150 mM aqueous NaCl, and simulated on the microsecond timescale. The lateral system size was at least 16 nm x 16 nm. All simulations were carried out at 310 K in the *NPT* ensemble using GROMACS versions 2021.7 and 2023.3 [86, 87].

**Table 3.** Compositions of the three plasma membrane models studied in this work. Numbers in parentheses indicate the number of cholesterol molecules in the respective leaflet at the end of the simulations. The CHOL(%) rows show the rounded percentage of cholesterol molecules in each leaflet. For lipid nomenclature see Figure 14.

| Name<br>$t_{\text{tot}}$ [$\mu\text{s}$] | Extracellular | | Cytoplasmic | | $N_{\text{ions}}$ | | $N_{\text{waters}}$ |
| --- | --- | --- | --- | --- | --- | --- | --- |
| | Lipid | $N_{\text{lipids}}$ | Lipid | $N_{\text{lipids}}$ | $\text{Na}^+$ | $\text{Cl}^-$ | |
| **ACD**<br>**20** $\mu\text{s}$ | LSM | 48 | OAPE | 32 | 271 | 151 | 62790 |
|  | NSM | 56 | PAPS | 120 |  |  |  |
|  | PSM | 72 | PDOPE | 72 |  |  |  |
|  | PAPC | 24 | PLQS | 80 |  |  |  |
|  | PLPC | 80 | PLPC | 80 |  |  |  |
|  | SOPC | 40 | POPC | 32 |  |  |  |
|  | CHOL | 404(401) | CHOL | 116(119) |  |  |  |
|  | CHOL(%) | 56% | CHOL(%) | 22% |  |  |  |
|  | PL<sub>all</sub> | 320 | PL<sub>all</sub> | 416 |  |  |  |
| **SCD**<br>**13** $\mu\text{s}$ | LSM | 48 | DPPC | 12 | 283 | 143 | 52134 |
|  | NSM | 56 | PAPS | 76 |  |  |  |
|  | PSM | 72 | PSM | 8 |  |  |  |
|  | PAPC | 32 | POPC | 20 |  |  |  |
|  | PLA20 | 16 | PLA20 | 68 |  |  |  |
|  | PLPC | 88 | PLPC | 52 |  |  |  |
|  | SAPS | 8 | SAPS | 8 |  |  |  |
|  | SOPC | 40 | POPE | 12 |  |  |  |
|  |  |  | PDOPE | 44 |  |  |  |
|  |  |  | SAPE | 20 |  |  |  |
|  |  |  | SAPI2A | 12 |  |  |  |
|  | CHOL | 240(243) | CHOL | 236(233) |  |  |  |
|  | CHOL(%) | 40% | CHOL(%) | 40% |  |  |  |
|  | PL<sub>all</sub> | 360 | PL<sub>all</sub> | 332 |  |  |  |
| **SCR**<br>**13** $\mu\text{s}$ | LSM | 24 | LSM | 24 | 278 | 138 | 58400 |
|  | NSM | 28 | NSM | 28 |  |  |  |
|  | PSM | 40 | PSM | 40 |  |  |  |
|  | PAPC | 16 | PAPC | 16 |  |  |  |
|  | PLA20 | 42 | PLA20 | 42 |  |  |  |
|  | PLPC | 70 | PLPC | 70 |  |  |  |
|  | SAPS | 8 | SAPS | 8 |  |  |  |
|  | SOPC | 20 | SOPC | 20 |  |  |  |
|  | DPPC | 6 | DPPC | 6 |  |  |  |
|  | PAPS | 38 | PAPS | 38 |  |  |  |
|  | PDOPE | 22 | PDOPE | 22 |  |  |  |
|  | POPC | 10 | POPC | 10 |  |  |  |
|  | POPE | 6 | POPE | 6 |  |  |  |
|  | SAPE | 10 | SAPE | 10 |  |  |  |
|  | SAPI2A | 6 | SAPI2A | 6 |  |  |  |
|  | CHOL | 238(241) | CHOL | 238(235) |  |  |  |
|  | CHOL(%) | 40% | CHOL(%) | 40% |  |  |  |
|  | PL<sub>all</sub> | 346 | PL<sub>all</sub> | 346 |  |  |  |

### System setup

Three PM models were investigated: a model with approximately **s**ymmetric lipid and **c**holesterol **d**ensities (SCD), a model with **a**symmetric lipid and **c**holesterol **d**ensities (ACD), and a **scr**ambled model (SCR). The SCD model was based on lipidomics and experimental measurements reported by Lorent *et al.* [9], whereas the ACD model followed the leaflet asymmetry proposed by Doktorova *et al.* [10]. Specifically, we adopted the leaflet-resolved lipid and cholesterol compositions reported by Doktorova *et al.*, which were derived from experimental constraints and refined through microsecond-timescale simulations [10]. The SCD membrane model and its interaction with the inhibitory Fc*γ*RIIb receptor was previously studied by our group [72]. Here, the protein was removed from the system, and the remaining lipid bilayer was simulated for 20 *µ*s in water with counterions only. To maintain consistency with the other systems studied here, the salt concentration was subsequently adjusted to 150 mM NaCl, and the system was simulated for an additional 13 *µ*s.

The main distinction between the SCD and ACD models was the degree of transbilayer phospholipid (PL) asymmetry. In the SCD model, the two leaflets contained comparable numbers of PL molecules. By contrast, in the ACD model, the cytoplasmic leaflet contained approximately 30% more PLs than the exoplasmic leaflet. This asymmetry is lower than the ∼50% imbalance estimated experimentally by Doktorova *et al.* [10]. To alleviate packing stress caused by leaflet area mismatch, cholesterol partitioned preferentially into the exoplasmic leaflet [10] (Table 3). The SCR model was generated from the SCD system by randomizing lipid distributions between leaflets, yielding symmetric lipid-type distributions while preserving the overall membrane composition. Each membrane contained approximately 40 mol% cholesterol in total. Following the same methodology, two additional bilayers with the same compositions and asymmetries as the ACD and SCD models, but with a quarter of the total number of lipids were simulated, referred to as sACD (15 *µ*s) and sSCD (1.7 *µ*s), respectively. The smaller bilayers were used to study the permeation of ethanol (EtOH) (see Materials and Methods, *Ethanol Simulations and Analysis*).

The ACD and SCR systems were generated using CHARMM-GUI [88–90]. For the ACD model, the plasmalogen PLQS (see Figure 14) was not available in the CHARMM-GUI lipid library. Therefore, PLA20 (see Figure 14) was used as a temporary substitute during bilayer assembly and solvation. These two lipids differ only in their *sn*-2 acyl chains: docosatetraenoic acid in PLQS (22:4) and arachidonic acid in PLA20 (20:4). After system assembly, PLA20 was replaced by PLQS using an in-house

**Fig. 14.**
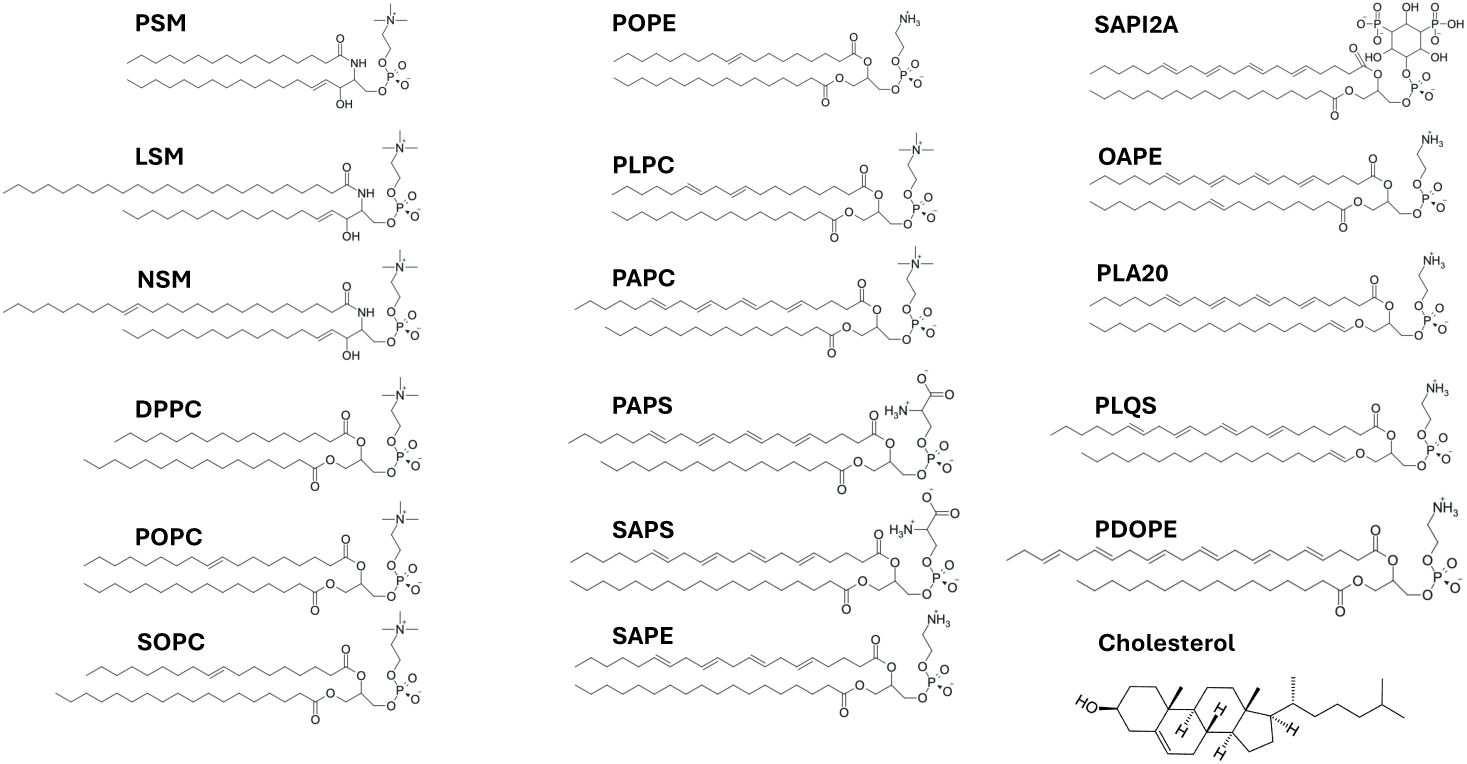
Chemical structures of lipid types used in the all-atom MD simulations.

Python script and newly generated PLQS topology files. All-atom simulations were performed using the GROMACS port of the CHARMM36 force field (July 2022) [91]. *General simulation parameters* For all simulations, the leap-frog integrator was used with a time step of 2 fs. Van der Waals interactions were described using a Lennard-Jones potential with a cut-off at 1.2 nm. A force-switch modifier was applied between 1.0 nm and 1.2 nm. Neighbor lists were handled using the Verlet scheme with a buffer radius of rlist=1.35 nm [92]. Long-range electrostatic interactions were computed via the fast smooth Particle-Mesh Ewald method [93, 94]. Temperature and pressure were maintained at 310 K and 1.0 bar, respectively. For production simulations, the v-rescale thermostat and the Parrinello-Rahman [95] or c-rescale [96] barostat were employed, with semi-isotropic coupling, a compressibility of 4.5 × 10*^−^*^5^ bar^-1^, and a time constant of *τ_P_* = 5.0 ps. Translational velocity of the center of mass was removed every 100 steps separately for the membrane and the solvent. Bonds involving hydrogens were converted to constraints propagated using the LINCS solver [97] with an order of 4 and one iteration.

System equilibration followed the protocol recommended by CHARMM-GUI, which consists of a single energy minimization followed by six short simulations with increasing time steps (1 fs to 2 fs) and varying levels of position/dihedral restraints on lipid headgroups and chiral centers. Final production simulations used for the comparison were run for 20 *µ*s (ACD), 13 *µ*s (SCD), and 13 *µ*s (SCR). The last 10 *µ*s were used for data collection.

### Ethanol simulations and analysis

A single ethanol molecule was inserted into the solvent phase of sACD and sSCD membranes. Membrane permeability was determined following the protocol of Wennberg et al. [98]. Briefly, the accelerated weight histogram (AWH) method [99, 100] was used to calculate the free energy profile, Δ*F* (*z*) (relative to bulk water), and the friction metric, *g*(*z*), as functions of position along the membrane normal, *z*, which served as the reaction coordinate. Following [98, 101], the ethanol permeability coefficient, *P*, was then calculated using the inhomogeneous solubility-diffusion model

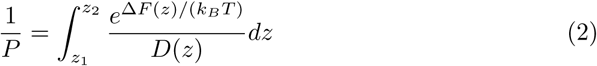

where *k_B_*is the Boltzmann constant, *T* is the temperature, and *D*(*z*) is the position-dependent diffusion coefficient. The integration was performed over the interval from −4 to 4 nm along the membrane normal. The diffusion coefficient was calculated as *D*(*z*) = *g^−^*^1^(*z*), after which a rolling median filter with a window width of 0.2 nm was applied to reduce noise in the data [98, 101]. The total simulation time was 11.7 *µ*s for the sSCD system and 10.3 *µ*s for the sACD system.

The convergence of the permeability coefficient, *P*, was monitored after both systems had passed the initial stage of the AWH simulations (Fig. 15a). The time evolution of *P* was fitted with an exponential decay function of the form *f* (*t*) = *Ae^−t/τ^* + *P*_inf_, where the fitted asymptote, *P*_inf_, was taken as the converged permeability coefficient.

**Fig. 15.**
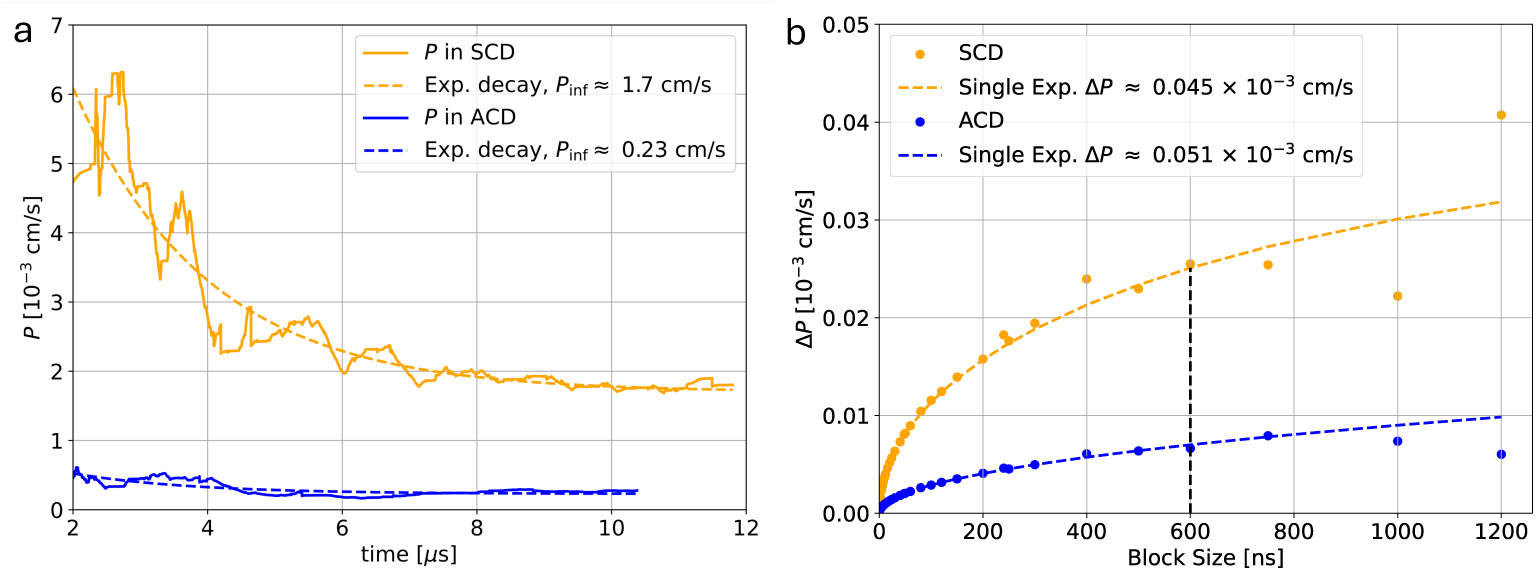
Permeability coefficient convergence and uncertainty. (a) Convergence of the permeability coefficient, *P*, over time for the sSCD (orange) and sACD (blue) systems. Dashed curves show fits to the function *f* (*t*) = *Ae^−t/τ^* + *P*_inf_. (b) Block averaging analysis of the *P* values obtained from the last 3 *µ*s of each simulation. Orange and blue dashed curves represent single-exponential fits following Hess [102] to the data below the black dashed line. The values reported in the legends correspond to the converged permeability coefficient, *P*_inf_, (left) and its estimated uncertainty, Δ*P*, (right) for each system.

To estimate the uncertainty, *P* values from the last 3 *µ*s of each simulation, during which the permeability coefficient had already converged, were used for block averaging (Figure 15b). The block-size dependence of the estimated error was fitted to a single-exponential function following Hess [102], yielding the final uncertainty estimate, Δ*P* (Figure 15b).

#### AWH Simulation Details

General simulation parameters were identical to those mentioned in Materials and Methods, *All-Atom Simulations*. Before each AWH simulation, a single ethanol molecule was inserted into the solvent phase of the corresponding system using *gmx insert-molecules* [86]. After a short energy minimization and equilibration with position restraints on the ethanol heavy atoms, the AWH simulations were initialized using the distance between the ethanol molecule and the membrane center of mass along the membrane normal as the single reaction coordinate. The reaction coordinate was sampled over the interval from −4 to 4 nm relative to the membrane center of mass. The initial diffusion constant along the reaction coordinate was set to 2 × 10*^−^*^5^ nm^2^ps*^−^*^1^, and a convolved biasing potential with a force constant of 25000 kJ mol*^−^*^1^nm*^−^*^2^ was applied.

### Cholesterol flip-flops

Cholesterol flip-flop (FF) events were defined as translocation of a cholesterol molecule between the outer and inner leaflets. FF analysis was performed using the *Lipyphilic* Python package [103]. In each frame, cholesterol molecules were assigned to leaflets based on the position of their hydroxyl oxygen atom (“O3” in CHARMM36 notation) relative to phospholipid phosphate atoms (“P” in CHARMM36 notation). Phospholipids did not undergo flip-flop on the simulated timescales.

A flip-flop attempt was recorded when the “O3” atom entered a 12 Å-wide slab centered at the membrane midplane. An attempt was classified as successful if the cholesterol reached the opposite leaflet and remained there for at least 10 ns; otherwise, it was classified as failed. The number of attempts was normalized by the average membrane area and analysis time to obtain a flip-flop rate (FFRate) for comparison between systems.

The local lipid environment during FF events was quantified using an enrichment index. Enrichment was measured from 35 ns to 10 ns before cholesterol’s hydroxyl oxygen atom changed leaflets and from 10 ns to 35 ns after the transition. The 10 ns intervals immediately before and after the transition were excluded because the enrichment index is ill-defined when cholesterol is near the membrane midplane and has not settled into either leaflet.

The instantaneous enrichment index of lipid species *B* around a transitioning cholesterol molecule, 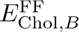, was defined as

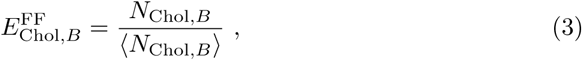

where *N*_Chol*,B*_ is the number of lipids of species *B* within 12 Å of the transitioning cholesterol, and ⟨*N*_Chol*,B*_⟩ is the average number of lipids of species *B* around any cholesterol molecule. Lipid species were grouped by acyl-chain unsaturation, defined as the total number of double bonds in their tails. For each unsaturation class, a weighted average enrichment was computed and reported in Table S1. For example, the enrichment of lipids with four or more unsaturations (“4U”) was calculated as

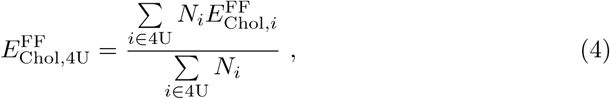

where *i* denotes a lipid species in the four-or-more-unsaturations class, *N_i_* is the number of lipids of species *i*, and 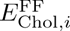 is the enrichment of species *i* around cholesterol.

### Area per lipid

The lipid-agnostic area per lipid (APL) was calculated by dividing the instantaneous simulation box area parallel to the membrane plane by the number of lipids (including cholesterol) per leaflet.

Lipid-type-specific APL was determined by 2D Voronoi tessellation of the instantaneous positions of phosphate atoms for PLs and hydroxyl oxygen atoms for cholesterol in each leaflet. The area *A* of each Voronoi polygon was calculated using the shoelace formula:

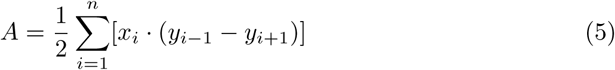

where (*x_i_, y_i_*) are the Cartesian coordinates of the *i*th vertex and *n* is the total number of vertices. Tessellation was performed using the *SciPy* package [104].

### *S*_CD_ order parameter

The deuterium order parameter *S*_CD_ was used to quantify phospholipid tail order, reflecting the average orientation of C–H bond vectors relative to the membrane normal. For tail carbon C*_i_*, *S*_CD_ was calculated as

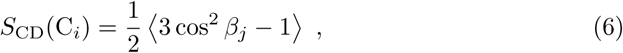

where *β_j_* is the angle between the membrane normal (here, the z-axis of the simulation system) and the C*_i_*–H*_j_* bond vector. The average was taken over all hydrogens on C*_i_*, over all equivalent carbons within the same lipid species, and over all analyzed frames.

### Lipid diffusion coefficients

In-plane diffusion coefficients (*D*) were calculated from the trajectories of lipid reference atoms: phosphate (“P”) for PLs and hydroxyl oxygen (“O3”) for cholesterol. Lipids were assigned to leaflets, and trajectories were unwrapped using the TOR scheme [105] to ensure continuity across periodic boundaries. Cholesterol molecules undergoing flip-flops (≈ 1%; Table S1) were excluded from the mean square displacement (MSD) analysis. Leaflet center-of-mass motion was removed independently for each leaflet.

The MSD for lipid *i* at lag time *τ* was defined as

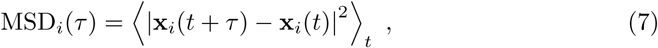

where ⟨·⟩*_t_* denotes an average over time origins. MSDs were averaged over lipids of the same species within each leaflet, and the diffusion coefficient was extracted by fitting the linear regime (*τ* ∈ [150 ns, 250 ns]), using

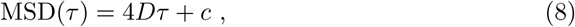

where *c* is a fitting constant. Uncertainties were estimated from the difference between diffusion coefficients determined from fits to the two halves of the full fitting interval, following the approach used in GROMACS’ *gmx msd* [86, 87].

### Local lipid environment

The local lipid environment was quantified using a nearest-neighbor enrichment index, *E*_AB_, based on 2D Voronoi tessellation of each leaflet. Phospholipids were represented by their central glycerol atom (“C2”/“C2S” in CHARMM36 notation) and cholesterol by the hydroxyl oxygen atom (“O3”). Nearest neighbors were defined as lipids sharing a Voronoi edge.

The enrichment of lipid species *B* around species *A* is defined as

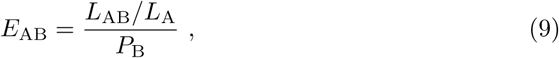

where

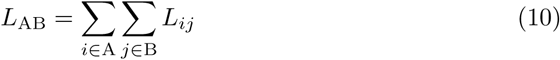

is the total Voronoi contact length shared between all lipids of species A and B. It is divided by

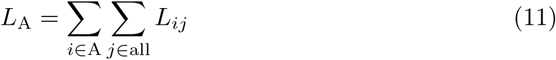

which is the total contact length of species *A* with all its neighbors. The reference contact probability of species B in a random mixture is

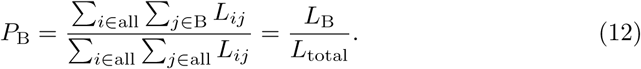

Thus, *L*_AB_*/L*_A_ is the fraction of species *A*’s contact length shared with species *B*, and *P*_B_ is the corresponding expectation for a random mixture, based only on the total Voronoi contact length contributed by species B. Values of *E*_AB_ *>* 1 indicate enrichment of species B around species A, whereas values of *E*_AB_ *<* 1 indicate depletion (Fig. 17).

**Fig. 16.**
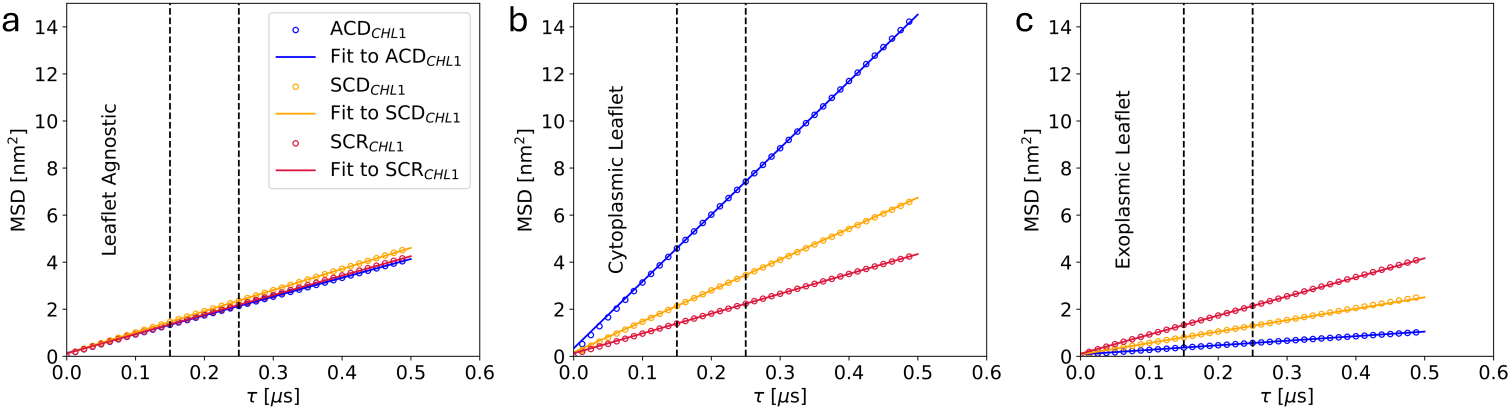
Mean square displacements (colored open circles) for cholesterol (CHL1) in the ACD (blue), SCD (orange), and SCR (crimson) systems. (a) Leaflet-agnostic MSDs; (b,c) MSDs for cholesterol molecules that remained exclusively in the cytoplasmic (b) or exoplasmic (c) leaflet. Solid lines are fits to Eq. 8; vertical dashed lines indicate the fitting interval.

**Fig. 17.**
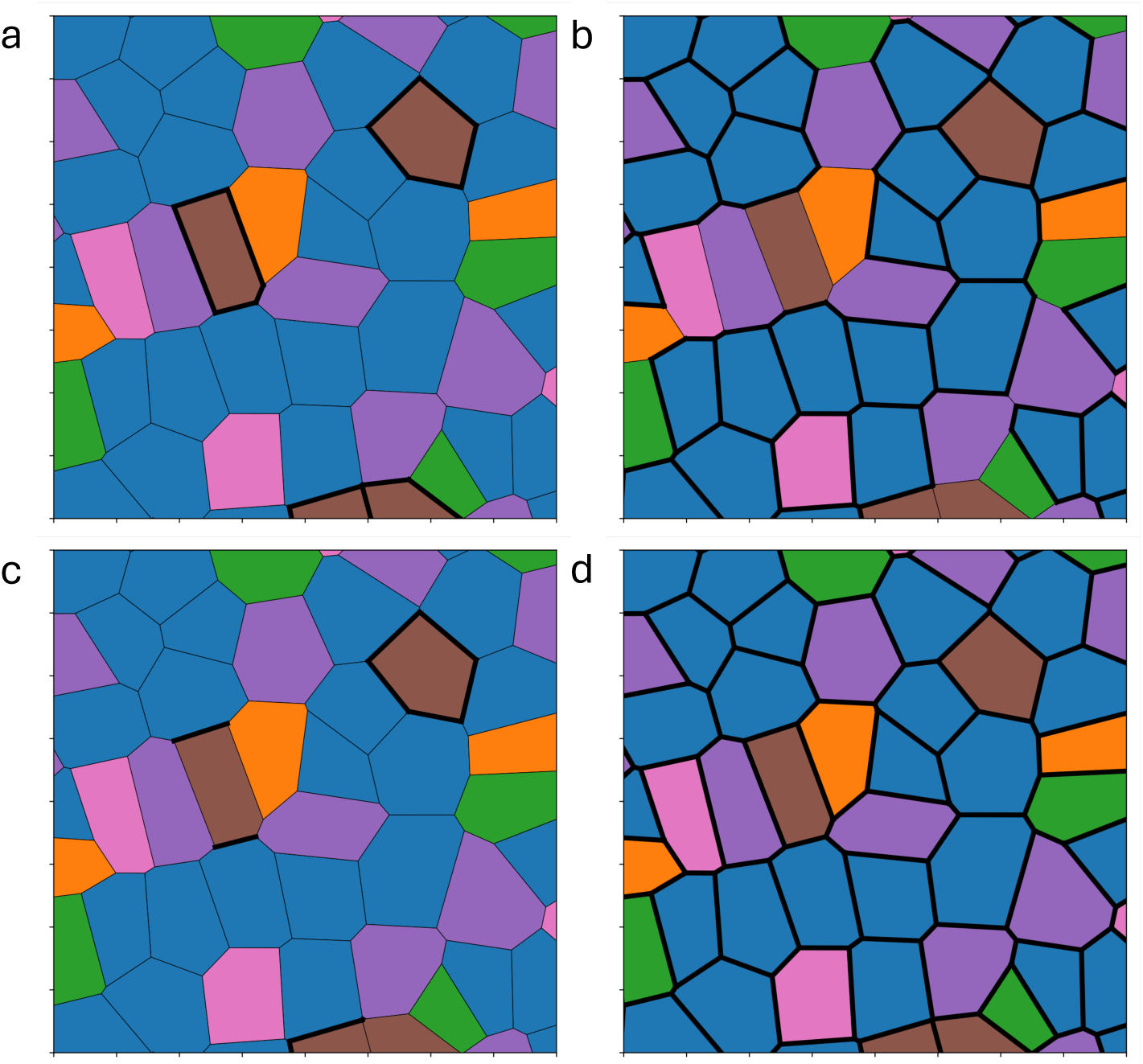
Schematic of the contact lengths used to compute the enrichment of species *B* (blue) around species *A* (brown). Thick black lines denote the contact edges included in *L*_A_ (a), *L*_B_ (b), *L*_AB_ (c), and *L*_total_ (d).

This edge-based definition avoids an arbitrary distance cutoff: unlike cutoff-based approaches, Voronoi neighbors share a direct contact edge, and weighting by edge length accounts for the extent of lipid–lipid contact.

### Area compressibility

The area compressibility modulus *K_a_* was calculated from simulation box area fluctuations [106–110]:

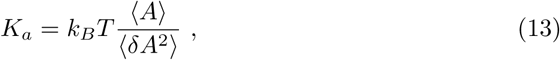

where *A* is the instantaneous membrane area projected onto the membrane plane, ⟨*A*⟩ is its mean, ⟨*δA*^2^⟩ = ⟨(*A* − ⟨*A*⟩)^2^⟩ is the mean-squared area fluctuation, *T* is the temperature, and *k_B_* is Boltzmann’s constant. Uncertainties were estimated by block-averaging [102].

### Bending

The membrane bending modulus *κ_b_* was calculated using the **L**ocal **F**luctuation **M**ethod (LFM) [32, 41].

*Grid assignment.* A scalar membrane height field *h_ij_* was constructed on a regular 2D grid (spacing ≈ 0.34 nm) using a modified version of the density mapping method of Ergüder et al. [111], which avoids empty grid cells. The height field was defined as the average of the upper- and lower-leaflet height fields:

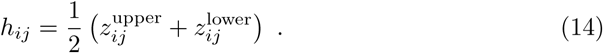

For each leaflet, 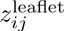 was the mean *z*-coordinate of all heavy atoms within a cylinder of radius *r_c_* = 8 Å centered at grid point (*i, j*), with *z* along the membrane normal. Leaflet assignments were updated at every analysis step to account for cholesterol flip-flops.

*Local fluctuation method*.

The LFM was applied to the height field as described in detail by Pöhnl *et al.* [32, 41]. Briefly, the instantaneous mean curvature *H̄* was calculated within circular patches of radius *r*. The probability distribution of *H̄* is

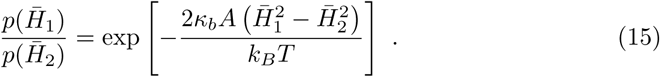

For an equilibrated membrane, *H̄* follows a Gaussian distribution with zero mean and variance *σ*^2^, so

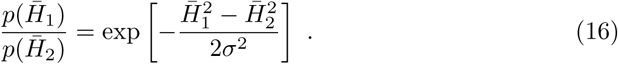

Combining Equations 15 and 16 yields the bending modulus:

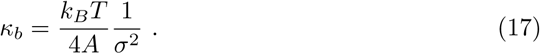

This value represents an upper bound for *κ_b_*, as lipid tilt is not explicitly included. The distribution was symmetrized to suppress artifacts from insufficient sampling. The procedure was repeated 1,000 times with randomly selected patch centers at fixed radius *r* = 30 Å, and the mean and uncertainty of *κ_b_* were estimated from the resulting distribution.

### Hydrophobic defects

#### 2D defects

Two-dimensional hydrophobic defect analysis was performed using PackMem [47]. The aim of this analysis is to identify membrane apolar regions that are vertically accessible to water. Lipids were assigned in each analyzed frame to the upper and lower leaflet, respectively. Lipid atoms were assigned van der Waals radii from the CHARMM36 force field [91]. Acyl-chain atoms below the carbonyl oxygens (or below the amide group in sphingomyelins) were classified as apolar; for cholesterol, all atoms except the hydroxyl-group were classified as apolar (Fig. 19a). Hydrophobic defects were defined as apolar atoms vertically exposed to the aqueous phase along the membrane normal. Shallow defects were located less than 1 Å below the central phospholipid glycerol atom; deep defects extended more than 1 Å below the same reference (Fig. 19b). Neighboring defects were clustered, and their combined area *A* was recorded. The resulting size distribution *p*(*A*) was fitted to

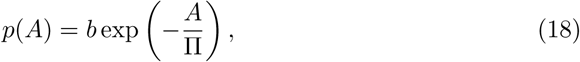

where *b* is a fitting constant and Π is the packing-defect constant.

**Fig. 18.**
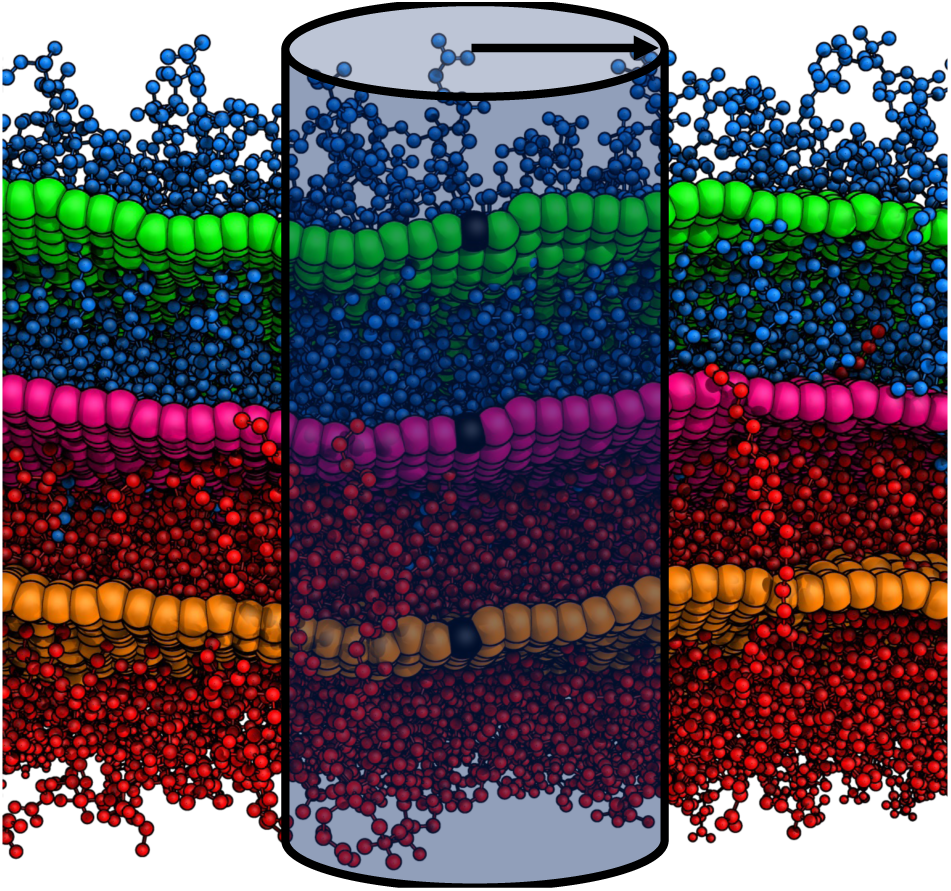
Height-field construction from a representative simulation frame. Heavy atoms in the upper and lower leaflets are shown in marine and red, respectively. For each grid point, the local upperleaflet height 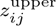 (green) and lower-leaflet height 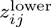 (orange) are computed from atoms within a cylinder of radius *r_c_*; their average gives the midplane height field *h_ij_* (hot pink) used for bending analysis. The cylinder radius is exaggerated for clarity.

#### 3D defects

Three-dimensional hydrophobic defects were identified using an algorithm similar to that of Tripathy *et al.* [48]:

1. A 3D grid with 1 Å resolution was generated to span the entire membrane.
2. The membrane excluded volume was determined as grid points not within the van der Waals radius of any lipid atom plus a probe radius *r_p_* = 1.4 Å (marine spheres in Fig. 19c,d).
3. A head-group grid was constructed from the positions of cholesterol hydroxyl oxygen atoms (“O3”) and the central glycerol atoms (“C2”/“C2S”) of phospholipids; grid points without a directly associated head-group atom were assigned values by nearest-neighbor interpolation (red spheres in Fig. 19c).
4. Excluded-volume grid points located between the upper and lower head-group grid were identified as defects (Fig. 19c,d).

**Fig. 19.**
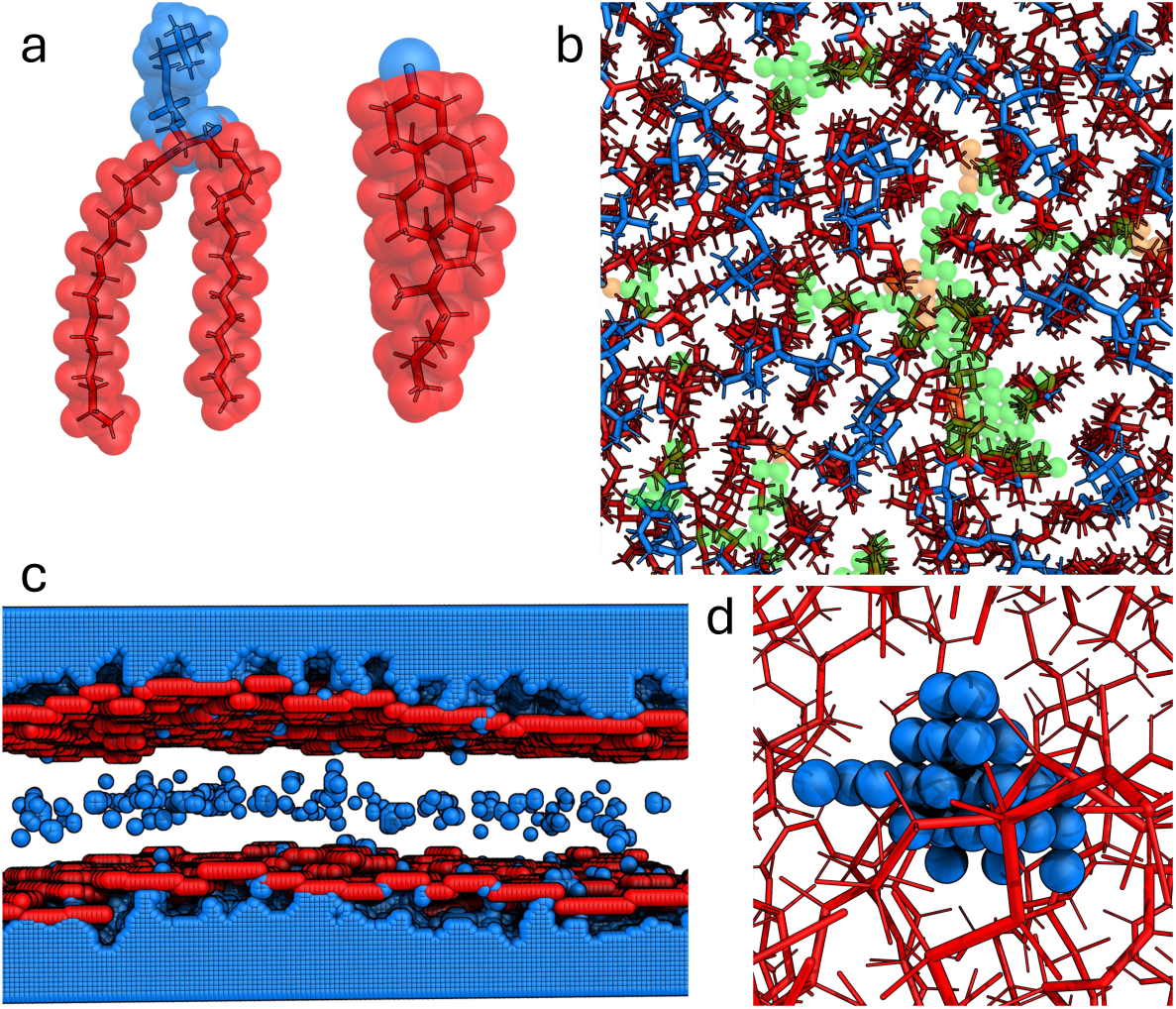
Identification of 2D (a,b) and 3D (c,d) hydrophobic defects. (a) Apolar (red) and polar (marine) atoms of a representative phospholipid and cholesterol molecule. (b) Deep (orange) and shallow (green) 2D defects. (c) Membrane excluded volume (marine spheres) and head-group grid (red spheres). (d) Zoomed-in view of a representative 3D defect (marine spheres) within the membrane interior (red sticks).

### Cholesterol clustering

Cholesterol clustering was analyzed by 2D Voronoi tessellation of each leaflet. Cholesterol molecules were represented by their hydroxyl oxygen atoms (“O3”) and phospholipids by their central glycerol atoms (“C2”/“C2S”). Nearest neighbors were defined as molecules sharing a Voronoi edge. A cholesterol molecule was assigned to a cluster if more than half of its Voronoi nearest neighbors were also cholesterol. Clusters were expanded iteratively by applying the same criterion to each nearestneighbor cholesterol of any cholesterol already assigned to the cluster, until no further additions were possible. Final clusters with less than 6 members were discarded.

### Defect Enrichment in cholesterol clusters

Cholesterol clusters and two-dimensional hydrophobic defects were identified independently for each leaflet (see Materials and Methods, *Hydrophobic Defects*). Defects were assigned to one of three populations according to the Voronoi cells in which they occurred: clustered cholesterol, nonclustered cholesterol, or phospholipids. For each population *i*, the total area *A^i^*was obtained from the corresponding Voronoi cells. The defect enrichment index was then calculated as

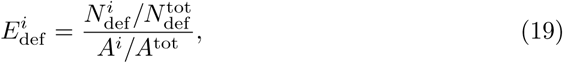

where 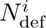 is the number of shallow or deep defects located within population *i*, 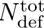 is the total number of defects in the leaflet, *A^i^* is the area occupied by population *i*, and *A*^tot^ is the total leaflet area. Thus, *E*_def_ = 1 corresponds to a defect density proportional to the area occupied by the population, whereas values greater or less than one indicate enrichment or depletion, respectively. Enrichment indices were calculated only for frames in which cholesterol clusters were present.

### Water permeability

#### Potential of mean force

On microsecond timescales, water molecules undergo multiple unbiased membrane permeation events, permitting estimation of the potential of mean force (PMF) along the membrane normal *z* from the equilibrium water density profile *ρ*(*z*):

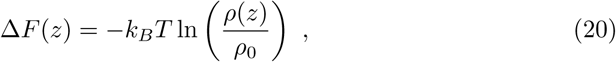

where *ρ*_0_ is the bulk water density (setting the bulk PMF to zero). Density profiles were computed with *gmx density* [86, 87].

#### Water permeation events

Water flux was calculated following Doktorova *et al.* [10]. A central slab of thickness 20 Å was defined around the bilayer midplane (−10 ≤ *z* ≤ 10 Å). Each water molecule was assigned one of three states per frame: +1 (above the slab), −1 (below), or 0 (inside). Permeation events were identified from the sequence of visited states: A successful permeation from the exoplasmic to the cytoplasmic side follows the sequence [+1, 0, −1], while the reverse follows [−1, 0, +1]. Unsuccessful attempts from the exoplasmic and cytoplasmic side are [+1, 0, +1] and [−1, 0, −1], respectively.

Transitions were not required to occur in consecutive frames; only the order of visited states was considered. Because water translocation events occur on timescales of hundreds of picoseconds or less, trajectories with a 10 ps coordinate output frequency were used. The total analyzed trajectory length was ≈ 1.2 *µ*s.

### Python usage

The programming language “Python” (python.org) was extensively used in this study, along with the modules Pandas [112], Seaborn [113], LiPyphilic [103], NumPy [114], MDAnalysis [115], SciPy [104], tqdm [116] and MatPlotLib [117].

## Data availability

All simulation models, input files and structures will be made available for download after acceptance of the manuscript.

## Acknowledgments

This work was supported by the German Research Foundation (Deutsche Forschungsgemeinschaft, DFG) through grants 534019561 and 555589280. The authors gratefully acknowledge the scientific support and HPC resources provided by the Erlangen National High Performance Computing Center (NHR@FAU) of the Friedrich-Alexander-Universität Erlangen-Nürnberg (FAU) under the NHR project IDs b223dc, b224dc, and b325dc. NHR funding is provided by federal and Bavarian state authorities.

## Author contributions

R.A.B. conceived and designed the study. C.R.P. performed the simulations. C.R.P., M.F.W.T., and C.Z. analyzed the data. C.R.P. and R.A.B. wrote the initial manuscript. All authors discussed the results, revised the manuscript, and approved the final version.

## Additional information

The supplementary information includes

- Supplementary Figures S1 – S4
- Supplementary Tables S1 – S4

## References

[1] Simons, K. & Ikonen, E. Functional rafts in cell membranes. Nature 387, 569–572 (1997).

[2] Wymann, M. P. & Schneiter, R. Lipid signalling in disease. Nat. Rev. Mol. Cell Bio. 9, 162–176 (2008).

[3] Hannun, Y. A. & Obeid, L. M. Principles of bioactive lipid signalling: lessons from sphingolipids. Nat. Rev. Mol. Cell Bio. 9, 139–150 (2008).

[4] Bretscher, M. S. Phosphatidyl-ethanolamine: differential labelling in intact cells and cell ghosts of human erythrocytes by a membrane-impermeable reagent. J. Mol. Biol. 71, 523–528 (1972).

[5] Verkleij, A. J. et al. The asymmetric distribution of phospholipids in the human red cell membrane. a combined study using phospholipases and freeze-etch electron microscopy. BBA Biomembranes 323, 178–193 (1973).

[6] Zwaal, R., Roelofsen, B., Comfurius, P. & Van Deenen, L. Organization of phospholipids in human red cell membranes as detected by the action of various purified phospholipases. BBA Biomembranes 406, 83–96 (1975).

[7] Van Meer, G., Gahmberg, C. G., Op den Kamp, J. A. F. & Van Deenen, L. L. M. Phospholipid distribution in human en(a-) red cell membranes which lack the major sialoglycoprotein, glycophorin a. FEBS Letters 135, 53–55 (1981).

[8] Murate, M. et al. Transbilayer distribution of lipids at nano scale. J. Cell Sci. 128, 1627–1638 (2015).

[9] Lorent, J. H. et al. Plasma membranes are asymmetric in lipid unsaturation, packing and protein shape. Nat. Chem. Biol. 16, 644–652 (2020).

[10] Doktorova, M. et al. Cell membranes sustain phospholipid imbalance via cholesterol asymmetry. Cell 188, 2586–2602.e24 (2025).

[11] Schütz, G. J. & Pabst, G. The asymmetric plasma membrane—a composite material combining different functionalities? balancing barrier function and fluidity for effective signaling. BioEssays 45, 2300116 (2023).

[12] Pabst, G. & Keller, S. Exploring membrane asymmetry and its effects on membrane proteins. Trends Biochem. Sci. 49, 333–345 (2024).

[13] Yamada, T. & Shinoda, W. Asymmetry and heterogeneity in the plasma membrane. Biophys. J. 125, 387–395 (2026).

[14] Doktorova, M., Symons, J. L. & Levental, I. Structural and functional consequences of reversible lipid asymmetry in living membranes. Nat. Chem. Biol. 16, 1321–1330 (2020).

[15] Wang, H.-Y. et al. Loss of lipid asymmetry facilitates plasma membrane blebbing by decreasing membrane lipid packing. Proc. Natl. Acad. Sci. U. S. A. 122, e2417145122 (2025).

[16] Nagata, S., Suzuki, J., Segawa, K. & Fujii, T. Exposure of phosphatidylserine on the cell surface. Cell Death & Differentiation 23, 952–961 (2016).

[17] Emoto, K., Inadome, H., Kanaho, Y., Narumiya, S. & Umeda, M. Local change in phospholipid composition at the cleavage furrow is essential for completion of cytokinesis. J. Biol. Chem. 280, 37901–37907 (2005).

[18] Sessions, A. & Horwitz, A. F. Myoblast aminophospholipid asymmetry differs from that of fibroblasts. FEBS Letters 134, 75–78 (1981).

[19] van den Eijnde, S. M., et al. Transient expression of phosphatidylserine at cell-cell contact areas is required for myotube formation. J. Cell Sci. 114, 3631–3642 (2001).

[20] Rival, C. M. et al. Phosphatidylserine on viable sperm and phagocytic machinery in oocytes regulate mammalian fertilization. Nat. Commun. 10, 4456 (2019).

[21] Emoto, K. et al. Redistribution of phosphatidylethanolamine at the cleavage furrow of dividing cells during cytokinesis. Proc. Natl. Acad. Sci. U. S. A. 93, 12867–12872 (1996).

[22] Emoto, K. & Umeda, M. An essential role for a membrane lipid in cytokinesis: regulation of contractile ring disassembly by redistribution of phosphatidylethanolamine. J. Cell Biol. 149, 1215–1224 (2000).

[23] Hammill, A. K., Uhr, J. W. & Scheuermann, R. H. Annexin v staining due to loss of membrane asymmetry can be reversible and precede commitment to apoptotic death. Exp. Cell Res. 251, 16–21 (1999).

[24] Ray, L. C. et al. Membrane association of monotopic phosphoglycosyl transferase underpins function. Nat. Chem. Biol. 14, 538–541 (2018).

[25] Kapus, A. & Janmey, P. Plasma membrane—cortical cytoskeleton interactions: A cell biology approach with biophysical considerations. Comprehensive Physiology 3, 1231–1281 (2013).

[26] Li, L., Shi, X., Guo, X., Li, H. & Xu, C. Ionic protein–lipid interaction at the plasma membrane: what can the charge do? Trends Biochem. Sci. 39, 130–140 (2014).

[27] Fisher, K. A. Analysis of membrane halves: cholesterol. Proc. Natl. Acad. Sci. U. S. A. 73, 173–177 (1976).

[28] Brasaemle, D. L., Robertson, A. D. & Attie, A. D. Transbilayer movement of cholesterol in the human erythrocyte membrane. J. Lipid Res. 29, 481–489 (1988).

[29] Schroeder, F. et al. Transmembrane distribution of sterol in the human erythrocyte. BBA Biomembranes 1066, 183–192 (1991).

[30] Lange, Y. & Slayton, J. M. Interaction of cholesterol and lysophosphatidylcholine in determining red cell shape. J. Lipid Res. 23, 1121–1127 (1982).

[31] Müller, P. & Herrmann, A. Rapid transbilayer movement of spin-labeled steroids in human erythrocytes and in liposomes. Biophys. J. 82, 1418–1428 (2002).

[32] Pöhnl, M., Kluge, C. & Böckmann, R. A. Lipid bicelles in the study of biomembrane characteristics. J. Chem. Theory Comput. 19, 1908–1921 (2023).

[33] Lange, Y. & Steck, T. L. Critique of a radically new model for plasma membrane bilayer organization. BioEssays 48, e70114 (2026).

[34] Skotland, T. & Sandvig, K. Plasma membranes: does one model fit all? Trends in Cell Biology (2026).

[35] Feigenson, G. W. Spiers memorial lecture: Experimental discovery of asymmetric bilayers, and a recent asymmetry example. Faraday Discussions 259, 9–25 (2025).

[36] Johnson, D. H., Kou, O. H., Bouzos, N. & Zeno, W. F. Protein–membrane interactions: sensing and generating curvature. Trends Biochem. Sci. 49, 401– 416 (2024).

[37] Khondker, A. et al. Membrane cholesterol reduces polymyxin b nephrotoxicity in renal membrane analogs. Biophys. J. 113, 2016–2028 (2017).

[38] Matsuzaki, K., Sugishita, K., Fujii, N. & Miyajima, K. Molecular basis for membrane selectivity of an antimicrobial peptide, magainin 2. Biochemistry 34, 3423–3429 (1995).

[39] Ruppelt, D. et al. The antimicrobial fibupeptide lugdunin forms water-filled channel structures in lipid membranes. Nat. Commun. 15, 3521 (2024).

[40] Rossetti, P. et al. From membrane composition to antimicrobial strategies: experimental and computational approaches to amp design and selectivity. Small 22, 2411476 (2026).

[41] Pöhnl, M., Trollmann, M. F. & Böckmann, R. A. Nonuniversal impact of cholesterol on membranes mobility, curvature sensing and elasticity. Nat. Commun. 14, 8038 (2023).

[42] Gu, R.-X., Baoukina, S. & Tieleman, D. P. Cholesterol flip-flop in heterogeneous membranes. J. Chem. Theory Comput. 15, 2064–2070 (2019).

[43] Bennett, W. D., MacCallum, J. L., Hinner, M. J., Marrink, S. J. & Tieleman, D. P. Molecular view of cholesterol flip-flop and chemical potential in different membrane environments. J. Am. Chem. Soc. 131, 12714–12720 (2009).

[44] Bennett, W. D. & Tieleman, D. P. Molecular simulation of rapid translocation of cholesterol, diacylglycerol, and ceramide in model raft and nonraft membranes. J. Lipid Res. 53, 421–429 (2012).

[45] Schrödinger, L. L. C. The PyMOL molecular graphics system, version 2.5.0 (2021).

[46] Róg, T. & Pasenkiewicz-Gierula, M. Cholesterol-sphingomyelin interactions: a molecular dynamics simulation study. Biophys. J. 91, 3756–3767 (2006).

[47] Gautier, R. et al. Packmem: a versatile tool to compute and visualize interfacial packing defects in lipid bilayers. Biophys. J. 115, 436–444 (2018).

[48] Tripathy, M., Thangamani, S. & Srivastava, A. Three-dimensional packing defects in lipid membrane as a function of membrane order. J. Chem. Theory Comput. 16, 7800–7816 (2020).

[49] Doktorova, M., LeVine, M. V., Khelashvili, G. & Weinstein, H. A new computational method for membrane compressibility: Bilayer mechanical thickness revisited. Biophys. J. 116, 487–502 (2019).

[50] Rawicz, W., Olbrich, K. C., McIntosh, T., Needham, D. & Evans, E. Effect of chain length and unsaturation on elasticity of lipid bilayers. Biophys. J. 79, 328–339 (2000).

[51] Rawicz, W., Smith, B., McIntosh, T., Simon, S. & Evans, E. Elasticity, strength, and water permeability of bilayers that contain raft microdomain-forming lipids. Biophys. J. 94, 4725–4736 (2008).

[52] Evans, E., Rawicz, W. & Smith, B. Concluding remarks back to the future: mechanics and thermodynamics of lipid biomembranes. Faraday discussions 161, 591–611 (2013).

[53] Kupferberg, J., Yokoyama, S. & Kezdy, F. The kinetics of the phospholipase a2catalyzed hydrolysis of egg phosphatidylcholine in unilamellar vesicles. product inhibition and its relief by serum albumin. J. Biol. Chem. 256, 6274–6281 (1981).

[54] Gordesky, S. E. & Marinetti, G. The asymmetric arrangement of phospholipids in the human erythrocyte membrane. Biochem. Biophys. Res. Commun. 50, 1027–1031 (1973).

[55] Tsamaloukas, A., Szadkowska, H. & Heerklotz, H. Thermodynamic comparison of the interactions of cholesterol with unsaturated phospholipid and sphingomyelins. Biophys. J. 90, 4479–4487 (2006).

[56] Engberg, O. et al. The affinity of cholesterol for different phospholipids affects lateral segregation in bilayers. Biophys. J. 111, 546–556 (2016).

[57] Slotte, J. P. Sphingomyelin–cholesterol interactions in biological and model membranes. Chem. Phys. Lipids 102, 13–27 (1999).

[58] Leventis, R. & Silvius, J. R. Use of cyclodextrins to monitor transbilayer movement and differential lipid affinities of cholesterol. Biophys. J. 81, 2257–2267 (2001).

[59] Levental, I., Levental, K. R. & Heberle, F. A. Lipid rafts: controversies resolved, mysteries remain. Trends in cell biology 30, 341–353 (2020).

[60] Mainali, L., Raguz, M. & Subczynski, W. K. Formation of cholesterol bilayer domains precedes formation of cholesterol crystals in cholesterol/dimyristoylphosphatidylcholine membranes: Epr and dsc studies. J. Phys. Chem. B 117, 8994–9003 (2013).

[61] Epand, R. M., Bach, D. & Wachtel, E. In vitro determination of the solubility limit of cholesterol in phospholipid bilayers. Chem. Phys. Lipids 199, 3–10 (2016).

[62] Stevens, M. M., Honerkamp-Smith, A. R. & Keller, S. L. Solubility limits of cholesterol, lanosterol, ergosterol, stigmasterol, and *β*-sitosterol in electroformed lipid vesicles. Soft matter 6, 5882–5890 (2010).

[63] Drin, G. et al. A general amphipathic *α*-helical motif for sensing membrane curvature. Nat. Struct. Mol. Biol. 14, 138–146 (2007).

[64] Bigay, J., Gounon, P., Robineau, S. & Antonny, B. Lipid packing sensed by arfgap1 couples copi coat disassembly to membrane bilayer curvature. Nature 426, 563–566 (2003).

[65] Pranke, I. M., et al. *α*-synuclein and alps motifs are membrane curvature sensors whose contrasting chemistry mediates selective vesicle binding. Journal of Cell Biology 194, 89–103 (2011).

[66] Bigay, J. & Antonny, B. Curvature, lipid packing, and electrostatics of membrane organelles: defining cellular territories in determining specificity. Developmental cell 23, 886–895 (2012).

[67] Vanni, S. et al. Amphipathic lipid packing sensor motifs: probing bilayer defects with hydrophobic residues. Biophys. J. 104, 575–584 (2013).

[68] Sikdar, S., Rani, G. & Vemparala, S. Role of lipid packing defects in determining membrane interactions of antimicrobial polymers. Langmuir 39, 4406–4412 (2023).

[69] Sato, Y. et al. Amphipathic helical peptide-nile red probes for fluorescence probing of the lipid packing defects and their surrounding membranes on exosomes. Sci. Rep. 15, 23790 (2025).

[70] Gierke, K. et al. The missing link: Piccolino is essential for tethering synaptic vesicles to rod photoreceptor ribbons. J. Cell Biol. 225, e202509110 (2026).

[71] Chuang, H.-L. et al. Visualizing dynamics of membrane rafts on live cells. Science Advances 11, eadv7001 (2025).

[72] Spiegel, F., et al. Role of lipid nanodomains for inhibitory fcriib function. bioRxiv (2023). URL https://www.biorxiv.org/content/early/2023/05/12/2023.05.09.540011.

[73] Head, B. P., Patel, H. H. & Insel, P. A. Interaction of membrane/lipid rafts with the cytoskeleton: impact on signaling and function: membrane/lipid rafts, mediators of cytoskeletal arrangement and cell signaling. BBA Biomembranes 1838, 532–545 (2014).

[74] Gómez-Llobregat, J., Buceta, J. & Reigada, R. Interplay of cytoskeletal activity and lipid phase stability in dynamic protein recruitment and clustering. Sci. Rep. 3, 2608 (2013).

[75] Chichili, G. R. & Rodgers, W. Cytoskeleton–membrane interactions in membrane raft structure. Cellular and molecular life sciences 66, 2319–2328 (2009).

[76] Heberle, F. A. & Doktorova, M. Exploring the sensitivities of experimental techniques to various types of membrane asymmetry using atomistic simulations. Faraday discussions 259, 300–320 (2025).

[77] Amaro, M., Reina, F., Hof, M., Eggeling, C. & Sezgin, E. Laurdan and di-4aneppdhq probe different properties of the membrane. Journal of physics D: Applied physics 50, 134004 (2017).

[78] Gupta, A., Korte, T., Herrmann, A. & Wohland, T. Plasma membrane asymmetry of lipid organization: fluorescence lifetime microscopy and correlation spectroscopy analysis. J. Lipid Res. 61, 252–266 (2020).

[79] Brahm, J. Permeability of human red cells to a homologous series of aliphatic alcohols. limitations of the continuous flow-tube method. J. Gen. Physiol. 81, 283–304 (1983).

[80] Evans, E. A., Waugh, R. & Melnik, L. Elastic area compressibility modulus of red cell membrane. Biophys. J. 16, 585–595 (1976).

[81] Waugh, R. & Evans, E. A. Thermoelasticity of red blood cell membrane. Biophys. J. 26, 115–131 (1979).

[82] Hossein, A. & Deserno, M. Spontaneous curvature, differential stress, and bending modulus of asymmetric lipid membranes. Biophys. J. 118, 624–642 (2020).

[83] Fiorin, G. & Forrest, L. R. Elucidating the mechanical properties of asymmetric membranes by direct derivation of their energetics. Faraday Discussions 259, 437–453 (2025).

[84] Brochard, F. & Lennon, J. Frequency spectrum of the flicker phenomenon in erythrocytes. Journal de Physique 36, 1035–1047 (1975).

[85] Evans, J., Gratzer, W., Mohandas, N., Parker, K. & Sleep, J. Fluctuations of the red blood cell membrane: relation to mechanical properties and lack of atp dependence. Biophys. J. 94, 4134–4144 (2008).

[86] Abraham, M. J. et al. Gromacs: High performance molecular simulations through multi-level parallelism from laptops to supercomputers. SoftwareX 1, 19–25 (2015).

[87] Páll, S., Abraham, M. J., Kutzner, C., Hess, B. & Lindahl, E. Tackling exascale software challenges in molecular dynamics simulations with gromacs, 3–27 (Springer, 2014).

[88] Jo, S., Kim, T., Iyer, V. G. & Im, W. Charmm-gui: a web-based graphical user interface for charmm. J. Comp. Chem. 29, 1859–1865 (2008).

[89] Wu, E. L., et al. Charmm-gui membrane builder toward realistic biological membrane simulations (2014).

[90] Brooks, B. R. et al. Charmm: the biomolecular simulation program. J. Comp. Chem. 30, 1545–1614 (2009).

[91] Klauda, J. B. et al. Update of the charmm all-atom additive force field for lipids: validation on six lipid types. J. Phys. Chem. B 114, 7830–7843 (2010).

[92] Kim, H., Fábián, B. & Hummer, G. Neighbor list artifacts in molecular dynamics simulations. J. Chem. Theory Comput. 19, 8919–8929 (2023).

[93] Darden, T., York, D. & Pedersen, L. Particle mesh ewald: An {N·log(N)} method for ewald sums in large systems. J. Chem. Phys. 98, 10089–10092 (1993).

[94] Essmann, U. et al. A smooth particle mesh ewald method. J. Chem. Phys. 103, 8577–8593 (1995).

[95] Parrinello, M. & Rahman, A. Polymorphic transitions in single crystals: A new molecular dynamics method. J. Appl. Phys. 52, 7182–7190 (1981).

[96] Bernetti, M. & Bussi, G. Pressure control using stochastic cell rescaling. J. Chem. Phys. 153, 114107 (2020).

[97] Hess, B., Bekker, H., Berendsen, H. J. C. & Fraaije, J. G. E. M. LINCS: A linear constraint solver for molecular simulations. J. Comput. Chem. 18, 1463–1472 (1997).

[98] Wennberg, C., Lundborg, M., Lindahl, E. & Norlen, L. Understanding drug skin permeation enhancers using molecular dynamics simulations. J. Chem. Inf. Model. 63, 4900–4911 (2023).

[99] Lidmar, J. Improving the efficiency of extended ensemble simulations: The accelerated weight histogram method. Physical Review E—Statistical, Nonlinear, and Soft Matter Physics 85, 056708 (2012).

[100] Lindahl, V., Lidmar, J. & Hess, B. Accelerated weight histogram method for exploring free energy landscapes. J. Chem. Phys. 141 (2014).

[101] Lundborg, M. et al. Skin permeability prediction with md simulation sampling spatial and alchemical reaction coordinates. Biophys. J. 121, 3837–3849 (2022).

[102] Hess, B. Determining the shear viscosity of model liquids from molecular dynamics simulations. J. Chem. Phys. 116, 209–217 (2002).

[103] Smith, P. & Lorenz, C. D. Lipyphilic: A python toolkit for the analysis of lipid membrane simulations. J. Chem. Theory Comput. 17, 5907–5919 (2021).

[104] Virtanen, P. et al. SciPy 1.0: Fundamental Algorithms for Scientific Computing in Python. Nature Methods 17, 261–272 (2020).

[105] Bullerjahn, J. T., von Bulow, S., Heidari, M., Henin, J. & Hummer, G. Unwrapping npt simulations to calculate diffusion coefficients. J. Chem. Theory Comput. 19, 3406–3417 (2023).

[106] Bochicchio, A., Brandner, A. F., Engberg, O., Huster, D. & Böckmann, R. A. Spontaneous membrane nanodomain formation in the absence or presence of the neurotransmitter serotonin. Frontiers in Cell and Developmental Biology 8, 601145 (2020).

[107] Helfrich, W. Elastic properties of lipid bilayers: theory and possible experiments. Zeitschrift für Naturforschung c 28, 693–703 (1973).

[108] Venable, R. M., Brown, F. L. & Pastor, R. W. Mechanical properties of lipid bilayers from molecular dynamics simulation. Chem. Phys. Lipids 192, 60–74 (2015).

[109] Feller, S. E. & Pastor, R. W. Constant surface tension simulations of lipid bilayers: the sensitivity of surface areas and compressibilities. J. Chem. Phys. 111, 1281–1287 (1999).

[110] Lindahl, E. & Edholm, O. Mesoscopic undulations and thickness fluctuations in lipid bilayers from molecular dynamics simulations. Biophys. J. 79, 426–433 (2000).

[111] Ergüder, M. F. & Deserno, M. Identifying systematic errors in a power spectral analysis of simulated lipid membranes. J. Chem. Phys. 154 (2021).

[112] pandas development team, T. pandas-dev/pandas: Pandas (2020). URL 10.5281/zenodo.3509134.

[113] Waskom, M. L. seaborn: statistical data visualization. Journal of Open Source Software 6, 3021 (2021). URL 10.21105/joss.03021.

[114] Harris, C. R. et al. Array programming with NumPy. Nature 585, 357–362 (2020). URL 10.1038/s41586-020-2649-2.

[115] Michaud-Agrawal, N., Denning, E. J., Woolf, T. B. & Beckstein, O. Mdanalysis: A toolkit for the analysis of molecular dynamics simulations. Journal of Computational Chemistry 32, 2319–2327 (2011). URL 10.1002/jcc.21787.

[116] da Costa-Luis, C. O. tqdm: A fast, extensible progress meter for python and cli. Journal of Open Source Software 4, 1277 (2019).

[117] Hunter, J. D. Matplotlib: A 2d graphics environment. Computing in Science & Engineering 9, 90–95 (2007).

