## Supplementary Information for "Large phospholipid-number asymmetry is not required to reproduce plasma membrane physical properties"

##### The PDF file includes

- Figures S1 – S4,
- Tables S1 – S4,
- References

### Supplementary Figures

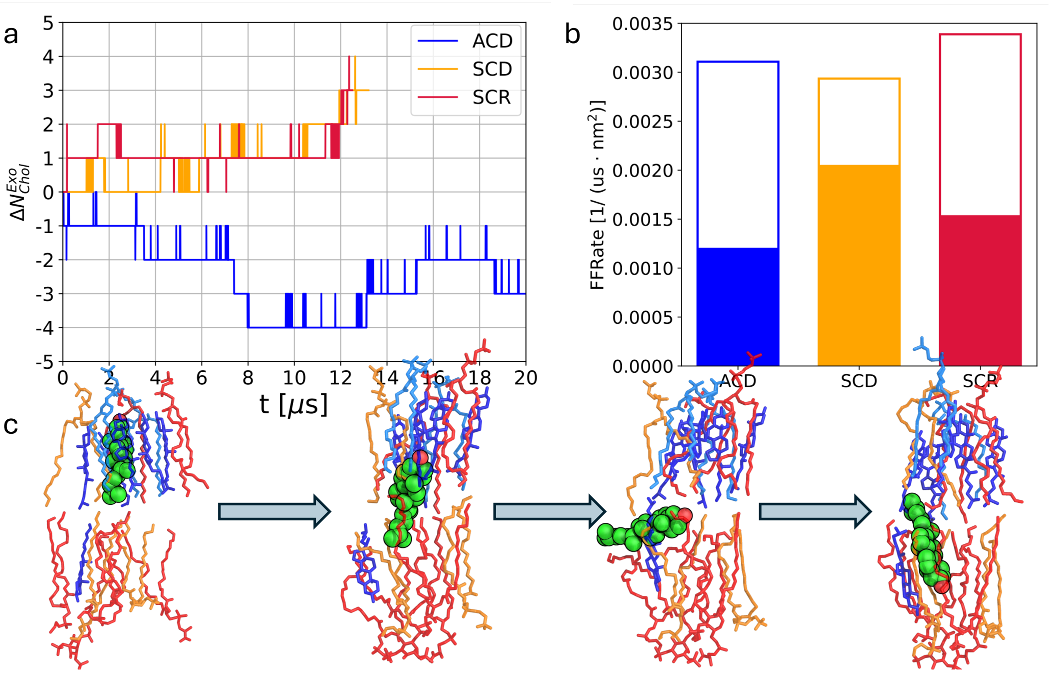


**Fig. S1 Cholesterol flip-flops.** (a) change in exoplasmic cholesterol number relative to the initial value, $\Delta N^{\mathrm{Exo}}_{\mathrm{Chol}}$. (b) rates of successful (filled bars) and unsuccessful (open bars) cholesterol flip-flop events. (c) representative successful flip-flop event, with the transitioning cholesterol shown as spheres and its local lipid environment shown as transparent sticks. Fully saturated lipids and sphingomyelins are shown in light blue, lipids with one or two unsaturations in orange, lipids with four or more unsaturations in red, and cholesterol in blue. Unsuccessful events similarly reach the membrane midplane but do not complete translocation to the opposite leaflet.

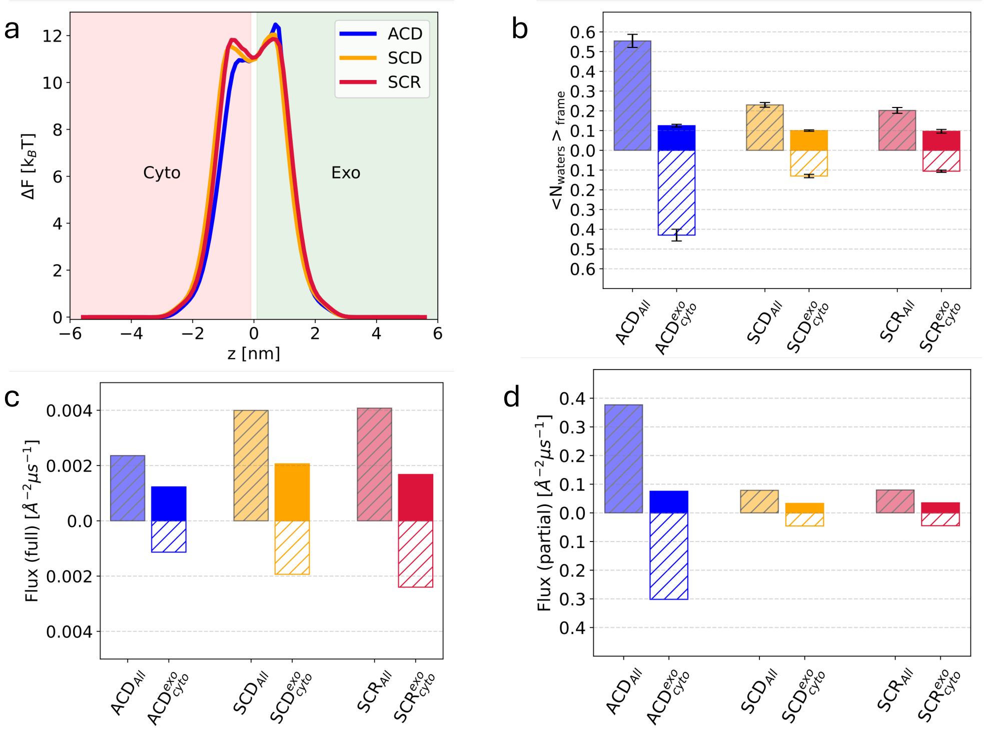


**Fig. S2 Water permeability.** (a) Potential of mean force (PMF), $\Delta F$, of water along the membrane normal for the three plasma membrane models. (b) Average number of water molecules per frame within the membrane midplane slab. Total values correspond to the full slab; filled and hatched components indicate the exoplasmic and cytoplasmic halves, respectively. (c) Successful water permeation flux, reported as the number of water molecules crossing the membrane per unit area and unit time. Total fluxes are shown together with exoplasmic-to-cytoplasmic and cytoplasmic-to-exoplasmic contributions. (d) Flux of partial permeation events, defined as water molecules entering the membrane midplane slab from one side and exiting again from the same side. The membrane midplane region was defined as a 20 Å slab centered at the bilayer midplane, extending 10 Å toward each leaflet.

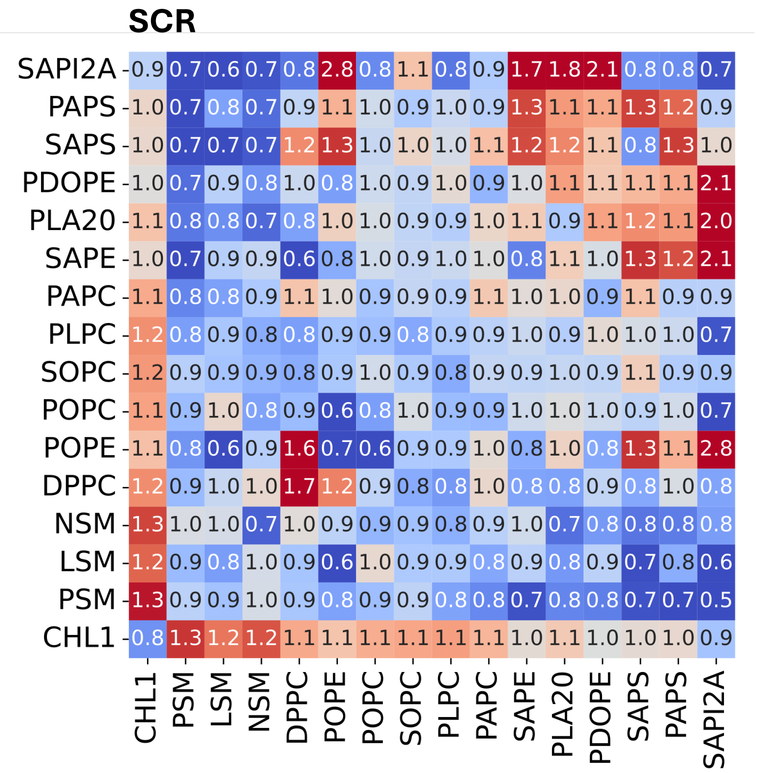


**Fig. S3 SCR enrichment matrix.** Lipid enrichment matrix for the symmetric SCR model. Enrichment indices were computed for each lipid species. The y-axis indicates the reference lipid type and the x-axis indicates the neighboring lipid type. Values greater than one indicate enrichment relative to a random mixture, whereas values below one indicate depletion. Lipids are ordered by degree of unsaturation and charge.

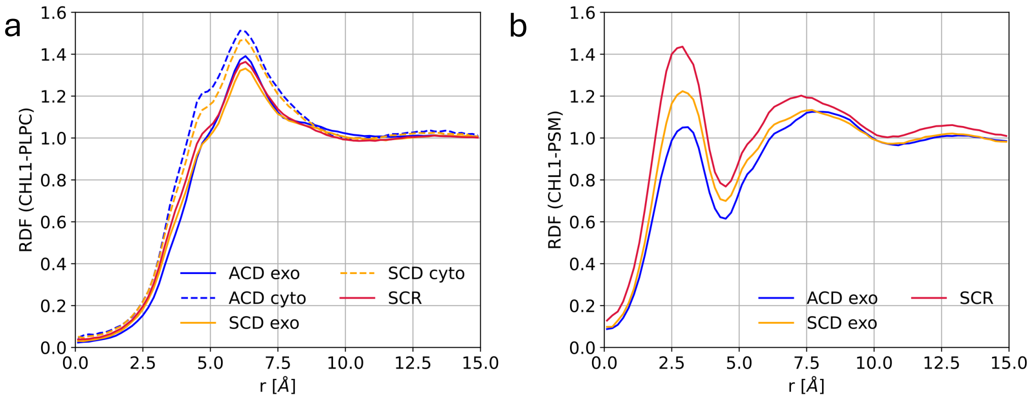


**Fig. S4 Radial distribution functions (RDFs) around cholesterol for representative phospholipid neighbors.** RDFs are shown for cholesterol–PLPC (a) and cholesterol–PSM (b) pairs. PLPC and PSM were selected because they are abundant and present in nearly all leaflets and systems.

#### Supplementary Tables

| **Sys-Ch. ID** | **Start LL** | **End LL** | $t_{\mathbf{trans}}$ [ns] | $\mathbf{E_{start}}$ | $\mathbf{E_{end}}$ |
|---|---|---|---|---|---|
| ACD-41 | Exo | Cyto | 29 | 1.9/1.3/**0**/0.6 | -/1.3/**0.9**/2.1 |
| ACD-172 | Exo | Cyto | 11 | 1.7/0.5/**2.4**/0.8 | -/0.6/**1.4**/0.7 |
| ACD-201 | Exo | Cyto | 10 | 0.8/0.9/**2.9**/0.8 | -/0/**1.3**/1.2 |
| ACD-235 | Exo | Cyto | 9 | 0.5/1.2/**3.1**/1.1 | -/0.8/**1.1**/1.6 |
| ACD-391 | Exo | Cyto | 52 | 0/1.7/**0**/0.7 | -/0.8/**1.1**/1.4 |
| ACD-458 | Cyto | Exo | 49 | -/0.9/**1.1**/1.2 | 0.7/1.3/**1.1**/1.0 |
| ACD-459 | Cyto | Exo | 16 | -/0.1/**1.6**/0.5 | 0.5/0.7/**3.4**/0.9 |
| SCD-6 | Exo | Cyto | 51 | 0.4/1.7/**1.2**/1.2 | 0/0.8/**1.4**/0.6 |
| SCD-73 | Cyto | Exo | 192 | 0.5/1.2/**1.3**/0.4 | 0.8/0.7/**2.2**/1.1 |
| SCD-102 | Cyto | Exo | 11 | 0/2.6/**0.9**/0.4 | 1.0/1.4/**1.2**/0.8 |
| SCD-186 | Cyto | Exo | 8 | 0/0.8/**1.3**/0.9 | 1.1/1.2/**1.5**/0.3 |
| SCD-199 | Cyto | Exo | 3 | 0/0.6/**1.2**/1.4 | 0.1/1.8/**2.5**/0.6 |
| SCD-359 | Exo | Cyto | 170 | 0.4/1.6/**1.6**/0.6 | 0/0.5/**1.6**/0.2 |
| SCD-474 | Cyto | Exo | 282 | 0/1.6/**1.0**/0.6 | 1.2/0.6/**0.2**/1.3 |
| SCR-97 | Exo | Cyto | 203 | 0.5/1.2/**1.7**/1.1 | 0.2/0.5/**2.1**/0.7 |
| SCR-279 | Cyto | Exo | 10 | 1.5/1.0/**1.2**/1.1 | 0.9/0.6/**1.5**/1.1 |
| SCR-360 | Cyto | Exo | 93 | 1.1/1.9/**0.5**/0.3 | 1.7/1.1/**0.2**/1.0 |
| SCR-433 | Cyto | Exo | 6 | 0.1/1.3/**2.0**/0.5 | 0.7/1.4/**1.6**/0.3 |
| SCR-471 | Cyto | Exo | 6 | 0.6/0.7/**1.5**/1.2 | 0.7/1.1/**2.1**/0.3 |

**Table S1** Successful cholesterol flip-flop events. The first column lists the system name (ACD, SCD, or SCR) and cholesterol identifier; the second and third columns indicate the initial and final leaflets; and the fourth column gives the transition time. The fifth and sixth columns report local lipid enrichment within 1.2 nm of the reference cholesterol before and after the event, respectively (see **Methods**). The final columns show enrichment grouped by lipid class: saturated, one- or two-unsaturated, **four-or-more-unsaturated**, and cholesterol. Values greater than one indicate local enrichment, whereas a dash denotes absence of the corresponding lipid class from the respective leaflet.

| Reference | Method | Compartment (d [nm]) | Reporter | D [$\mu m^2/s$] |
|---|---|---|---|---|
| Tank *et al.* [1] | FPR | CS L6 Myoblast | NBD-PC | 0.44 ± 0.15 |
| | FPR | Bleb L6 Myoblast | NBD-PC | 1.2 ± 0.4 |
| | FPR | CS FDB | NBD-PC | 0.1 ± 0.05 |
| | FPR | Bleb FBD | NBD-PC | 1.5 ± 0.6 |
| Chai *et al.* [2] | COBRI-M | CS PtK2 (¡100 nm) | DOPE-PEG2000-biotin + AuNP | $0.49_{slow}$ / $0.84_{fast}$ |
| | COBRI-M | CS PtK2 (¡100 nm) | DSPE-PEG2000-biotin + AuNP | $0.47_{slow}$ / $0.86_{fast}$ |
| Andrade *et al.* [3] | STED-FCS | CS NRKF (40 nm) | DPPE + Atto647N | 0.56 ± 0.04 |
| | STED-FCS | CS $MEFs_{IA32}$ (40 nm) | DPPE + Atto647N | 0.72 ± 0.04 |
| Doktorova *et al.* [4] | FCS | $MEFs_{3T3}$ | TF-SM | 1.25 ± 0.17 |
| | FCS | $MEFs_{3T3}$ | TF-Chol | 2.0 ± 0.24 |
| | FCS | $MEFs_{3T3}$ Scr | TF-SM | 2.07 ± 0.10 |
| | FCS | $MEFs_{3T3}$ Scr | TF-Chol | 2.44 ± 0.33 |
| Current work | AA-MD | $ACD_{exo}$ | SM | 0.39 ± 0.00 |
| | AA-MD | $ACD_{exo}$ | Chol | 0.49 ± 0.00 |
| | AA-MD | $SCD_{exo}$ | SM | 0.95 ± 0.01 |
| | AA-MD | $SCD_{exo}$ | Chol | 1.22 ± 0.02 |
| | AA-MD | SCR | SM | 1.52 ± 0.03 |
| | AA-MD | SCR | Chol | 2.03 ± 0.03 |

**Table S2** Diffusion coefficients ($D$) summarized. d denotes the observation diameter where specified. FPR, fluorescence photobleaching recovery; NBD-PC, N-4-nitrobenz-2-oxa-1,3-diazole phosphatidylcholine; CS, cell surface; FDB, flexor digitorum brevis muscle fibers; COBRI-M, contrast-enhanced coherent brightfield microscopy; PtK2, epithelial kidney cells from long-nosed potoroo; DO(S)PE-PEG2000-biotin, PEGylated and biotinylated DO(S)PE lipid; AuNP, gold nanoparticle (≈ 30 nm diameter); STED-FCS, stimulated emission depletion fluorescence correlation spectroscopy; NRKF, normal rat kidney fibroblasts; MEF, mouse embryonic fibroblasts; DPPE-Atto647N, phosphatidylethanolamine labeled with a fluorescent dye at the headgroup; TF-SM, TopFluor sphingomyelin (dipyrrometheneboron difluoride-labeled); TF-Chol, TopFluor cholesterol; Scr, ionophore-induced scrambling; AA-MD, all-atom molecular dynamics simulations. $D_{slow}$ and $D_{fast}$ denote distinct diffusion modes reported for the corresponding probes.

| Reference | Method | Membrane | Temperature [K] | Bending Modulus [$k_B$T] |
|---|---|---|---|---|
| Méléard *et al.* [5] | Flicker Analysis | Total RBC LE | 310.15 | 34.1 ± 3.7 |
| | Flicker Analysis | Total RBC LE | 298.15 | 65.1 ± 4.6 |
| | Flicker Analysis | Total RBC LE | 298.15 | 86.5 ± 3.4 |
| | Flicker Analysis | Total RBC LE | 298.15 | 114.2 ± 4.9 |
| Graciá *et al.* [6] | Flicker Analysis | PM RBC $LE_1$ | 296.15 | 55.0 ± 2.0 |
| | Flicker Analysis | PM RBC $LE_2$ | 296.15 | 55.0 ± 3.2 |
| Himbert *et al.* [7] | Diffuse X-ray Scattering | RBC ghosts | 310.15 | 2-6 |
| | Neutron Spin-Echo | RBC ghosts | 310.15 | 4-7 |
| | Coarse Grained MD | RBC based on [8, 9] | 310.15 | 4 |
| Current Work | AA MD | ACD | 310.00 | 38.1 ± 0.1 |
| | AA MD | SCD | 310.00 | 36.6 ± 0.1 |
| | AA MD | SCR | 310.00 | 36.2 ± 0.1 |

**Table S3** Summary of bending moduli. RBC LE - red blood cell lipid extracts. Total RBC LE - RBC LE of the whole cell with resulting lipid composition PC:PE:SM:PS:Chol = 20.4:16.2:14.4:9.0:40. PM RBC $LE_1$ - RBC LE only from the plasma membrane of a RBC, with composition - PC:PE:SM:PS:Chol = 14:14:11:10:50. PM RBC $LE_2$ - similar as $LE_1$ but with 40% cholesterol and containing transmembrane peptides. Note that RBC leaflet asymmetry is lost during lipid extract preparation [10]. For comparison the total membrane composition for SCR and SCD is PC:PE:SM:PS:Chol ≈ 21:13:16:9:41; ACD - PC:PE:SM:PS:Chol ≈ 20:15:14:10:41.

| **Reference / method** | **SM** | **PC** | **PE** | **PS** |
|---|---|---|---|---|
| Verkleij *et al.* [11], *Naja naja* $PLA_2$ | 82 | 76 | 20 | 0 |
| van Meer *et al.* [12], *Naja naja* $PLA_2$ | 78 | 80 | 17 | 2 |
| Zwaal *et al.* [13], *Naja naja* $PLA_2$ | 85 | 68 | 0 | 0 |
| Zwaal *et al.* [13], *Apis mellifera* $PLA_2$ | 85 | 55 | 9 | 0 |
| Lorent *et al.* [14], *Apis mellifera* $PLA_2$ | 89 | 48 | 4 | 15 |
| Doktorova *et al.* [4], *Apis mellifera* $PLA_2$ | 90 | 45 | 0 | 0 |
| Murate *et al.* [15], SDS-FRL | 98.5 | 98.1 | 0 | 0.7 |
| Bretscher [16], FMMP | – | – | 0 | 0 |
| Gordesky and Marinetti [17], TNBS | – | – | 30 | 0 |

**Table S4** Reported exoplasmic fractions of major phospholipid classes in human red blood cell membranes. Values indicate the percentage of a given lipid class assigned to the exoplasmic leaflet. Where applicable, the origin of $PLA_2$ is indicated. SM, sphingomyelin; PC, phosphatidylcholine; PE, phosphatidylethanolamine; PS, phosphatidylserine; SDS-FRL, sodium dodecyl sulfate-digested freeze-fracture replica labeling; FMMP, formyl methionyl sulfone methyl phosphate labeling; TNBS, trinitrobenzene sulfonic acid labeling. Dashes indicate that the respective lipid class was not analyzed.

### References


[1] Tank, D. W., Wu, E.-S. & Webb, W. W. Enhanced molecular diffusibility in muscle membrane blebs: release of lateral constraints. *The Journal of cell biology* **92**, 207–212 (1982).

[2] Chai, Y.-J., Cheng, C.-Y., Liao, Y.-H., Lin, C.-H. & Hsieh, C.-L. Heterogeneous nanoscopic lipid diffusion in the live cell membrane and its dependency on cholesterol. *Biophysical Journal* **121**, 3146–3161 (2022).

[3] Andrade, D. M. *et al.* Cortical actin networks induce spatio-temporal confinement of phospholipids in the plasma membrane–a minimally invasive investigation by sted-fcs. *Scientific reports* **5**, 11454 (2015).

[4] Doktorova, M. *et al.* Cell membranes sustain phospholipid imbalance via cholesterol asymmetry. *Cell* **188**, 2586–2602.e24 (2025).

[5] Meleard, P. *et al.* Bending elasticities of model membranes: influences of temperature and sterol content. *Biophysical journal* **72**, 2616–2629 (1997).

[6] Gracia, R. S., Bezlyepkina, N., Knorr, R. L., Lipowsky, R. & Dimova, R. Effect of cholesterol on the rigidity of saturated and unsaturated membranes: fluctuation and electrodeformation analysis of giant vesicles. *Soft Matter* **6**, 1472–1482 (2010).

[7] Himbert, S. *et al.* The bending rigidity of the red blood cell cytoplasmic membrane. *PLoS One* **17**, e0269619 (2022).

[8] Stefanoni, D. *et al.* Red blood cell metabolism in rhesus macaques and humans: comparative biology of blood storage. *Haematologica* **105**, 2174 (2019).

[9] Phillips, G. B. & Dodge, J. T. Composition of phospholipids and of phospholipid fatty acids of human plasma. *Journal of lipid research* **8**, 676–681 (1967).

[10] Bligh, E. G. & Dyer, W. J. A rapid method of total lipid extraction and purification. *Canadian journal of biochemistry and physiology* **37**, 911–917 (1959).

[11] Verkleij, A. J. *et al.* The asymmetric distribution of phospholipids in the human red cell membrane. a combined study using phospholipases and freeze-etch electron microscopy. *Biochimica et Biophysica Acta (BBA)-Biomembranes* **323**, 178–193 (1973).

[12] Van Meer, G., Gahmberg, C. G., van Deenen, L. *et al.* Phospholipid distribution in human en (a-) red cell membranes which lack the major sialoglycoprotein, glycophorin a (1981).

[13] Zwaal, R., Roelofsen, B., Comfurius, P. & Van Deenen, L. Organization of phospholipids in human red cell membranes as detected by the action of various purified phospholipases. *Biochimica et Biophysica Acta (BBA)-Biomembranes*

**406**, 83–96 (1975).

[14] Lorent, J. H. *et al.* Plasma membranes are asymmetric in lipid unsaturation, packing and protein shape. *Nat. Chem. Biol.* **16**, 644–652 (2020).

[15] Murate, M. *et al.* Transbilayer distribution of lipids at nano scale. *Journal of cell science* **128**, 1627–1638 (2015).

[16] Bretscher, M. S. Phosphatidyl-ethanolamine: differential labelling in intact cells and cell ghosts of human erythrocytes by a membrane-impermeable reagent. *Journal of molecular biology* **71**, 523–528 (1972).

[17] Gordesky, S. E. & Marinetti, G. The asymmetric arrangement of phospholipids in the human erythrocyte membrane. *Biochemical and Biophysical Research Communications* **50**, 1027–1031 (1973).
